# Neuropeptide F regulates adult female response to diet, larval locomotion and several larval physiological processes in *Lucilia cuprina cuprina*

**DOI:** 10.64898/2025.12.22.696103

**Authors:** Juan P. Wulff, Esther J. Belikoff, Vanesa A. S. Cunha, Pedro Mariano-Martins, Maxwell J. Scott

## Abstract

The blowfly *Lucilia cuprina dorsalis* is a highly detrimental ectoparasite of sheep responsible for causing flystrike, a condition that can result in severe morbidity and potentially lead to death if left untreated. In contrast, *Lucilia cuprina cuprina* is necrophagous and not a pest. We have been interested in identifying the genes that may play a role in the evolution of a parasitic lifestyle in blowflies. The objective of the present work was to explore the physiological role of the *L. cuprina long neuropeptide F* gene (*LcNPF*) in *L. c. cuprina.* We used CRISPR/Ca9 to generate a strain carrying a loss-of-function knock-in mutation for *LcNPF*. An RNA-Seq analysis was performed, and physiological and behavioral assays were conducted to evaluate the role of *LcNPF* in larvae and adult flies. Our findings indicate that functional disruption of the *LcNPF* gene significantly impairs egg hatching, larval survival, weight gain, crawling speed and larval time to pupariation. These phenotypic changes were corroborated by RNA-Seq analysis, which revealed downregulation of transcripts in *LcNPF* null mutated larvae associated with the respective physiological processes. In contrast, the foraging behavior of the larvae under the tested conditions was not affected. Interestingly, NPF appears to be essential for the oviposition preference of females for rotten meat but not for male mating behavior or fertility. Our results suggest that NPF signaling plays a central regulatory role in multiple physiological processes across both the larval and adult stages.

**Highlights:**

□ A loss-of-function knock-in mutation of the *LcNPF* gene was obtained using CRISPR/Cas9.
□ *LcNPF*-homozygous mutant larvae showed decreased weight gain, locomotion and survival on rotten meat.
□ Disruption of the *LcNPF* gene eliminated the female preference for oviposition in rotten meat.
□ *LcNPF* does not appear to be essential for normal fertility or male mating behavior as homozygotes were comparable to wild type

## 1. Introduction

The Australian sheep-blowfly *Lucilia cuprina* Wiedemann, 1830 (Diptera: Calliphoridae) can exhibit both parasitic and necrophagous lifestyles and these ecological strategies vary across geographic regions and subspecies. The subspecies *Lucilia cuprina dorsalis*, which occur across sub-Saharan Africa and the Australasian region, is predominantly parasitic and is a major pest of sheep (Doll, 2020; Kapoor et al., 2025b, 2025a, 2024; Waterhouse and Paramonovo, 1950). Conventional insecticides are the current control method of *L. c. dorsalis* (Heath and Levot, 2015); however, resistance to the use of insecticides such as organophosphates and benzoylphenyl urea has been reported (Heath and Levot, 2015). Conversely, the subspecies *Lucilia cuprina cuprina* is predominately necrophagous and is more widely distributed globally, including Asia, Australia, and the Americas (Kapoor et al., 2024; Nelson et al., 2012). In the New World, *L. c. cuprina* is necrophagous, developing mainly on decomposing matter and animal remains, with only rare reports of myiasis in humans (Doll, 2020; Kapoor et al., 2025a; Mouga and Gaedke, 2017; Sanford, 2017; Waterhouse and Paramonovo, 1950).

The *L. cuprina* substrate seeking behavior for oviposition has been extensively studied (Ashworth and Wall, 1994). The main attractants to gravid females are sulfur-rich compounds, ammonia and other volatiles emitted from decomposing organic material, such as meat or feces (Ashworth and Wall, 1994; Benbow et al., 2015; Forbes and Perrault, 2014). Dimethyl disulfide (DMDS), dimethyl trisulfide (DMTS), Indole and p-cresol are among the most relevant blowfly attractants emitted by decomposing organic material (Chaudhury et al., 2010; Liu et al., 2016; Zhu et al., 2013). In this regard, it has been observed that the last larval stage and adult gravid females of *L. c. cuprina* are more attracted to 5-day-old rotten beef compared to fresh beef (Cunha et al., 2023; Wulff et al., 2024).

*Lucilia c. dorsalis* females lay eggs in open wounds or moist tissues of host animals, and the hatched larvae consume the animal tissue producing large wounds that can lead to death (Byrd and Tomberlin, 2019; Fenton et al., 1999). The immature larval stages congregate in clusters (Fenton et al., 1999) and this arrangement enhances the action of larval secretions and excretions (SE). Larval SE includes peptidases, lipases, ammonia and antimicrobial peptides among the most relevant components (Casu et al., 1994; Hobson RP, 1932; Lennox, 1940). In addition, SE are associated with exogenous predigestion of animal tissue and larval immune response to host defenses and biological agents (such as bacteria and fungi), allowing the larvae to feed, survive and complete their immature stages (Ashworth and Wall, 1994).

Neuropeptides and their receptors (G protein-coupled receptors, GPCRs) play a critical integrative role regulating all physiological processes and behaviors in insects, such as feeding, immune response and oviposition (Schoofs et al., 2017). Consequently, they have been suggested as potential targets for pest control (Fónagy, 2006; Scherkenbeck and Zdobinsky, 2009). Neuropeptides have been well studied in blowflies including, short and long neuropeptides F, FMRFamide, Myosuppressin, Sulfakinin, leucokinins, Allatostatin, Adipokinetic Hormone (AKH), insulin-like peptides (ILPs), Tachykinin, and the neuropeptide Pigment Dispersing Factor (PDF) (Angioy et al., 2007; Downer et al., 2007; Duve et al., 1979; Duve and Thorpe, 1994; Gäde et al., 1990; Haselton et al., 2006; Matsushima et al., 2003; Setzu et al., 2012). However, these studies were restricted to the blowflies *Phormia regina* (Meigen, 1826), *Protophormia terraenovae* Robineau-Desvoidy, 1830, and *Calliphora vomitoria* (Linnaeus, 1758), and mostly focused on feeding behavior and tissue contraction, among a few other physiological processes (Angioy et al., 2007; Downer et al., 2007; Duve et al., 1979).

In insects, the long neuropeptide F (NPF) is initially produced as a prepropeptide with an amino terminal signal peptide that facilitates secretion, which is removed during translation (CUI and ZHAO, 2020). The propeptide contains a dibasic cleavage site that is recognized and cleaved by enzymes to produce the mature NPF peptide typically ranging from 36 to 40 amino acids in most species (CUI and ZHAO, 2020). The carboxyl terminus is amidated and characterized by the motif RXRFamide (CUI and ZHAO, 2020; Fadda et al., 2019). NPF occurs predominantly in nervous tissue and gut, and to a lesser extent in other organs such as the fat body and mouthparts (CUI and ZHAO, 2020). NPF has been linked to various physiological processes throughout different insect species, including ovary development (Cerstiaensa et al., 1999), feeding and sleep-wake behaviors (Chung et al., 2017; Williams et al., 2020), courtship behavior and reproduction (Liu et al., 2019; Van Wielendaele et al., 2013), immune response (Deng and Chiu, 2022) and cardiac activity (Setzu et al., 2012) among other functions (CUI and ZHAO, 2020). To date, only the role on cardiac activity has been investigated in a blowfly species (Setzu et al., 2012). Furthermore, when assessing the larval feeding behavior of the blowfly *Cochliomyia hominivorax* (Coquerel, 1858), reared on rotten meat, NPF expression was higher in individuals that avoided this substrate, suggesting that this gene may influence behavioral or physiological processes associated with rotten meat (Amaral Faria, 2019).

The main objective of the present work was to explore the physiological role of NPF in *L. c. cuprina.* To this end, we generated an *L. c. cuprina* homozygous strain carrying a loss-of-function mutation for *LcNPF*. Following, an RNA-Seq analysis was performed on single larvae at 3 and 5 days after egg hatching, that were heterozygous or homozygous for a *LcNPF* null mutation. Furthermore, physiological and behavioral assays were conducted to evaluate the role of *LcNPF* in larvae and adult flies.

## 2. Materials and Methods

### 2.1. Individuals used for experiments

All larvae and flies used in the present study were of the sub-species *L. c. cuprina*, called “*L. cuprina*” to ease the reading of the manuscript. Wild-type (*wt*) flies belonged to the *L. cuprina*-LA07 *wt* colony, which was established in our laboratory in 2010 using approximately 300 pupae of mixed sexes provided by Dr. Aaron Tarone (Texas A&M, TX, USA). The LA07 colony was established by Dr. Tarone through multiple adult-fly collections in Los Angeles, CA, USA in 2007. In addition, larval and adult heterozygous and homozygous for the *LcNPF* knock-in strain were used for the RNA-Seq analysis and behavioral assays. All larvae and adult flies used for the experiments were reared using protocols previously described (Li et al., 2014), kept at 23.5 ± 1 °C and a non-controlled photoperiod (∼13:11 light/dark).

### 2.2. Generation of *Lucilia cuprina* NPF loss-of-function mutation

#### 2.2.1. *LcNPF* gene, guide RNA (gRNA) and CRISPR/Cas9 in vitro assay

The *LcNPF* genomic and proteomic sequences, detailing the guide (g) RNA used for gene editing, including the PAM sequence and the Cas9 cutting site, are shown in the supplementary method (SM) section SM1A-C. The *LcNPF* gene model was plotted using SnapGene (Dotmatics, Boston, MA, USA) and edited with Adobe Illustrator (Adobe, San Jose, CA, USA). In addition, the cutting efficiency of the gRNA selected for the experiments, *i.e*. LcupNPF-gRNA-11, was tested using a Cas9 in vitro assay, which is described at SM2A-B.

#### 2.2.2. DNA donor construction, CRISPR/Cas9 mix preparation and eggs injections

The construction of the *LcNPF* homology direct repair (HDR) dsDNA donor is described in detail at SM3A-E. Briefly, an *LcNPF* gene fragment corresponding to the left (L) and right (R) homology arms (HA) of the dsDNA donor was amplified by polymerase chain reaction (PCR). The purified PCR product was ligated with the pUC19 plasmid, and the resulting plasmid was subsequently linearized by restriction digestion and ligated with a synthetic DNA fragment that included NotI and XhoI restriction sites. To obtain the final construct, the plasmid including the gene HAs was digested with NotI and XhoI and ligated with a *Lchsp83-ZsGreen*-*tub3’* fragment that was excised from the donor plasmid pB[LcHsp83ZsGreenTubpA] (Concha et al., 2011), using the same enzymes. The sequence of the final dsDNA donor plasmid (to be used for injections) was confirmed by Oxford Nanopore whole plasmid DNA sequencing.

Protocols used for CRISPR/Cas9 mixture, needles preparation and embryo injections were the same previously described by (Wulff et al., 2025). Briefly, Cas9 (IDT, Cat. # 1081058) was mixed with the duplex crRNA::tracrRNA, 1M Potassium chloride (KCl) and 1 μl of 3.1 buffer (NEB, Cat. # B6003S). The LcNPF knockin plasmid was added to the mix to reach a final plasmid concentration of 500 ng/μl. Final concentrations for the other compounds were as follows: Cas9 = 3.1 μM (500 ng/μl); crRNA::tracrRNA = 7.5 μM (250 ng/μl) and KCl = 250 mM.

The *LcNPF* gene model generated for Fig. 1 showing the gene editing site was obtained with SnapGene (Dotmatics) and edited with Adobe Illustrator (Adobe). The LcNPF protein secondary structure was plotted with Protter (https://wlab.ethz.ch/protter/start/), and figure sections arranged with BioRender (bioRender, Toronto, Ontario, Canada).

**Figure 1.**
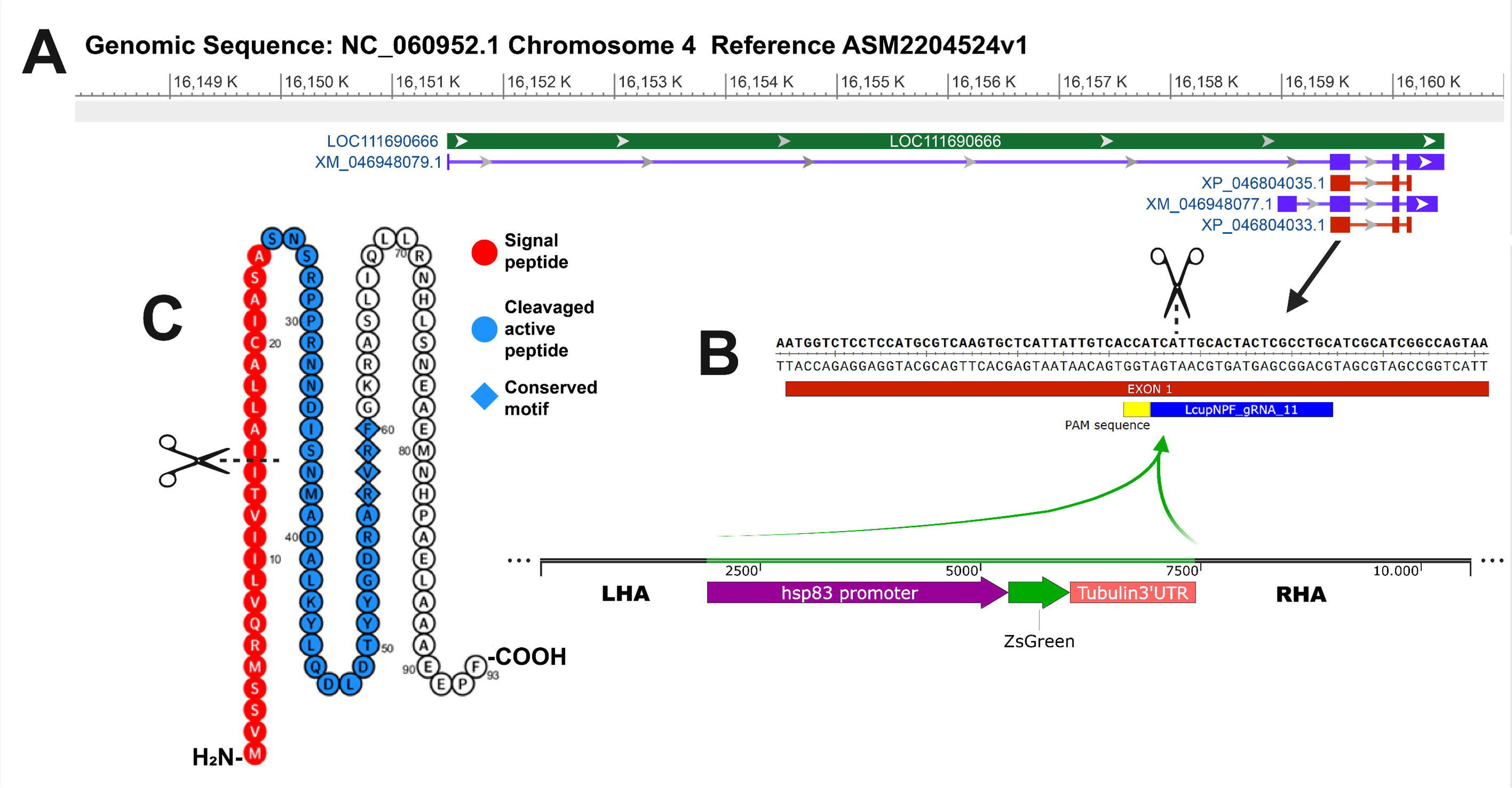
CRISPR/Cas9-mediated loss-of-function mutation of the *L. cuprina* NPF gene (*LcNPF*). **A**. Shows the locus, transcripts and CDS of the *LcNPF* gene, detailed in green, purple and red respectively. **B**. Exon 1 gene sequence showing Cas9 cutting site, gRNA and PAM sequence. The *Lchsp83-ZsGreen* transgene flanked by the right (R) and left (L) homology arms (HA) are shown below. **C**. Protein sequence highlighting the signal peptide, active peptide, conserved motif and site of the interruption of the protein sequence (scissors). For more details about *LcNPF* knockin pipeline refer to Figs. S1A-J.

#### 2.2.3. Fly screening, crosses and genotyping

Larvae that hatched from injected eggs were screened using a Leica M205 FA microscope with the ZsGreen filter set (EX: 500/20; EM: 535/30). Only mosaic larvae of generation G0 showing transient expression of the plasmid carrying the ZsGreen marker were retained to obtain adult flies (SM4A). Emerged adult flies were crossed following protocols described in (Wulff et al., 2025). After two backcrosses vs. *wt* flies, followed by a sibling cross of heterozygous (HET) individuals (SM4B), homozygous third-instar larvae (L3) were identified based on the intensity of green fluorescence (SM4C) and bred to establish the *LcNPF^-/-^* mutant strain.

To confirm the *Lchsp83-ZsGreen* gene had inserted into the *LcNPF* gene at the correct location, whole single *LcNPF^-/-^* larvae were placed in 2.0 mL tubes prefilled with zirconium beads (Benchmark Scientific, Tempe, AZ, USA, Cat. #D1032-30) and 500 µl of DNA extraction buffer (Biosearch Technologies, Hoddesdon, UK, Cat. #QE09050). Larvae were disrupted for 30 s using a benchtop homogenizer (Benchmark Scientific, Cat. #Z742475) set to 6.5 m/s. Subsequently the tubes were incubated for 20 min at 65°C and 2 min at 98°C. Following, tubes were vortexed for 5 s and centrifuged for ∼10 s at 14000 x g at room temperature (RT). The whole homogenate was transferred to a clean 1.5 mL Eppendorf tube, and the DNA was purified using the DNA Clean Kit (Zymo Research, Irvine, CA, USA, Cat. #D4004) following manufacturer’s specifications. The genomic DNA was used as a template for PCR using primers flanking the *LcNPF* gene, and from within the transgene. The PCR products were then cloned, sequenced and blasted to the *L. cuprina* genome assembly ASM2204524v1 as described in SM5A-B.

### 2.3. RNA-Seq experiment

#### 2.3.1. Sample collection and RNA isolation

To complete the RNA-Seq analysis, a *LcNPF^+/-^* heterozygous (HET) or *LcNPF^-/-^* homozygous (HOM) single whole larva was removed from the meat diet at different days after egg hatching: day 3 (D3), which included a mix of late second (late-L2) and early third (early-L3) larval instar; and day 5 (D5), including only late third larval instar (late-L3). The HET larvae were obtained by crossing *LcNPF^-/-^* homozygotes females to wild type males. The HOM cultures were set at the same time and reared under identical conditions. Samples were arranged into four groups, namely, HETD3, HOMD3, HETD5 and HOMD5 (n = 4 groups), including five samples (replicates) per group.

Larvae were recovered from rearing buckets and placed in 1.5 mL tubes filled with 300 µl of cold RNAlater^TM^ (Thermo-Fisher, Waltham, MA, USA, Cat. #AM7020) and stored at −80 °C until use. Prior to RNA extraction, the RNAlater^TM^ was removed, and larvae rinsed two times with 1 mL of cold 50% ethanol. After rinsing, larvae were resuspended in 600 µl of cold Trizol^TM^ and homogenized using pellet pestles (Bel-Art, Wayne, NJ, USA, Cat. #F65000-0002) and a homogenizer (Bel-Art, Cat. #F65100-0000) for 90 s. Subsequently, samples were centrifuged at 16000 x g and 4 °C for 5 min to pellet part of lipids, cuticle and other debris. After centrifugation the supernatant was transferred to a clean 1.5 mL tube. Total RNA was extracted using the Quick-RNA^TM^ kit (Zymo Research, Cat. #R1050) according to the manufacturer’s specifications. The exception was the addition of a second deoxyribonuclease 1 (DNase 1) step to ensure the absence of genomic DNA contamination in the samples, as follows: 1) during the RNA extraction 30 U of DNAse1 plus 75 µl of digestion buffer (DB) were added to the crude samples followed by 15 min of incubation at room temperature (RT); 2) after elution of the clean RNA in nuclease-free (NF) water, another 5 U of DNAse1 + 2.5 µl of DB were added followed by an incubation time same as above. After DNAse1 treatments, the RNA Clean & Concentrator^TM^-5 kit (Zymo Research, Cat. #R1013), was used to purify the RNA following the manufacturer’s specifications. The RNA was eluted in 12 μl of nuclease free water, quantified using a Qubit-4^TM^ fluorometer with the HS kit (Thermo-Fisher, Cat. # Q33120) and kept at −80°C until use.

#### 2.3.2. RNA-Seq data analysis

Sample quality control, library construction and sequencing services were provided by Novogene Inc. (Sacramento, CA, USA), following protocols described by the provider (https://www.novogene.com/us-en/resources/downloads/). The sequencing platform was Illumina NovaSeq 6000 (Illumina, San Diego, CA, USA), using paired end 150 bp reads and the sequencing depth coverage was ∼50 million raw reads per library. The Geneious Prime® software v2023.2.1 (https://www.geneious.com) and associated plugin packages were used to complete all the bioinformatic analysis and generate the RNA-Seq data output. BBDuk plugin v38.88 was utilized to remove Illumina adaptors and trim off low-quality bases at the 5′ and 3′ends using the following parameters: kmer lengh: 27; trim both ends: minimum quality Q20; trim adapters based on paired reads overhangs: minimum overlap 22; discard short reads: minimum length 20 bp. Trimmed reads were mapped to the reference *L. cuprina* genome assembly NCBI ID ASM2204524v1 using the default Geneious Prime mapper set to detect all types of RNA sequences and a low-medium sensitivity. Gene expression was calculated only for identified coding sequence (CDS) and the normalization method used was transcript per million (TPM), but RPKMs (Reads Per Kilobase per Million mapped reads) and FPKMs (Fragments Per Kilobase per Million mapped fragments) were also provided with the raw data.

Differential gene expressions between groups were calculated using DESeq2 (Love et al., 2014). An FDR Benjamini-Hochberg (BH) adjusted *P*-value < 0.05 and Log_₂_ fold-change (Log_₂_FC) > 1 was used to define differentially expressed (DE) transcripts between groups. For the differential gene expression analysis, the comparisons were completed between HOM and HET larvae, at each developmental time. Days (instead of larval stages) were chosen to define compared groups because day 3 included a mix of L2 and L3 larval stages. In addition, no comparisons were made between samples collected on different days. The objective was to identify down and upregulated genes in the homozygous larvae (NPF null mutated) relative to heterozygous larvae, at each chosen developmental time (always expressed throughout this manuscript as days post-hatching to enhance clarity and readability). Further, a principal components analysis (PCA) was performed using Geneious Prime® software and the DESeq2 normalization method, to compare all libraries.

### 2.4. *LcNPF^-/-^* strain fitness evaluation

#### 2.4.1. Fertility

To determine the fertility of *wt* and *LcNPF^-/-^*flies, a single *wt* or NPF null mutant fly (*LcNPF^-/-^*) was crossed with two *wt* flies of the opposite sex, using protocols adapted from (Concha et al., 2016). For oviposition, females were offered fresh 93/7% ground beef overnight (∼15 h). A total of 25 single crosses per condition were completed.

#### 2.4.2. Eggs hatching rate and tolerance to desiccation

Six-hundred eggs were obtained from 10-day-old gravid females of the LA07 *wt* and *LcNPF^-/-^* colonies (300 eggs per colony), using fresh 93/7% ground beef as oviposition substrate. Hundred eggs of each genotype were separated, prepared, desiccated and incubated following the same protocol described for uninjected eggs in section: *DNA donor construction, CRISPR/Cas9 mix preparation and embryo (egg) injections.* The other 200 eggs of each genotype were processed using the same protocol except for the desiccation step. Eggs were screened after 19-26 h to record egg hatching.

#### 2.4.3. Weight of L2 and L3 larvae reared using lean-fat 73/27 and 93/7% ground beef

Lean-fat 73/27% or 93/7% ground beef was offered as oviposition substrate to 10-day-old gravid females of the LA07 *wt* and *LcNPF^-/-^*colonies. Eggs were let reach an early L2 stage on the same substrate following previously described protocols (Li et al., 2014). To avoid overcrowded clusters of larvae (associated with decreased larval weight and survival), hundred and fifty mg of early L2 larvae (∼250 larvae; 1 day after egg hatching), were transferred onto fresh beef of the same type previously used as the oviposition substrate. This larval counting method was selected due to the difficulty of accurately quantifying this high number of small larvae necessary for all replicates. The approximate number of larvae per unit-weight was previously estimated by counting and weighing 10 groups of 50 larvae. In addition, 30 larvae of each genotype and reared using each type of beef were separated and weighed at this developmental time. Weight of larvae was measured at 2-, 3- and 6-day-old (after egg hatching), early and late L2 and late L3, respectively. For each type of beef, the sample size was n = 30 larvae for both L2 stages and n = 60 for the L3 stage.

#### 2.4.4. Larval locomotion

Locomotion activity of wandering late-L3 larvae was recorded, *i.e*. time vs. distance when crawling in a straight line using a protocol adapted from (Post and Paululat, 2018). Larvae were obtained from the LA07 *wt* and *LcNPF^-/-^* colonies and reared in lean-fat fresh 93/7% ground beef at room temperature (Li et al., 2014). Wandering late-L3 stage larvae were filmed while crawling in an arena to calculate the covered distance vs. time (SM6). No stimulus was used to lure the larvae. Only larvae exhibiting straight-line crawling (Video S1) were selected for analysis until counting n = 25 larvae per compared group.

#### 2.4.5. Larval survival assay using beef under different conditions

The effect on larvae survival, number of larvae that reach the pupal stage and pupal emergence rate, was evaluated using different types of beef divided into two assays: 1) larvae were reared using fresh or rotten lean-fat 73/27% or 93/7% ground beef. The rotten beef was obtained by placing each type of fresh beef at 30°C and 50% relative humidity for 5 days; and 2) larvae were reared using fresh 93/7% ground beef alone, supplemented with ammonium hydroxide (Fisher Scientific, Waltham, MA, USA, Cat. #A669500) or supplemented with urea (Sigma-Aldrich, Cat. # U5128-100G) to reach a 1% final concentration for both chemicals. The doses selected for ammonia and urea were defined after observing that higher concentrations were harmful to *wt* larvae. To assess survival on each type of diet, the LA07 *wt* and *LcNPF^-/-^*colonies were offered fresh 93/7% ground beef to collect eggs (Li et al., 2014). Subsequently, eggs were let to hatch and reach an early-L2 stage (1 day after egg hatching). At this point, hundred-and-fifty mg of early L2 larvae (∼250 larvae) were transferred to each type of beef and reared to reach adulthood. Individuals were screened at days 3, 5, 7, 9, 11, 15 and 25 after egg hatching, to record number of dead larvae, pupae and adult emergence. Three replicates per type of diet were conducted for each genotype of larvae, *i.e*. LA07 *wt* and *LcupNPF^-/-^*. Comparisons were made between different types of beef within each genotype.

#### 2.4.6. Larval pupariation

Larvae were reared in 93/7% ground beef (Li et al., 2014). A total of n = 100 L3 wandering larvae (6 days after egg hatching) was isolated from the LA07 *wt* and *LcNPF^-/-^* colonies. The number of pupae for each group was recorded at 7, 8, 9 and 10 days after egg hatching.

### 2.5. Behavioral assays

*LcNPF*^-/-^ and *wt* larval or adult individuals were used in each of the following sections.

#### 2.5.1. Larval diet preference test

The larval diet preference test was performed following protocols described previously (Cunha et al., 2023). About 120 3-day-old early L3 larvae were used for the assays. The choices offered to the larvae were rotten (R) and fresh (F) 93/7% ground beef, either at 25 ± 1 or 33 ± 1°C, designated as cold (C) and hot (H) respectively. Those larvae that did not choose any option after 10 min were considered as non-choice (NC). For further details about beef conditions refer to (Cunha et al., 2023).

#### 2.5.2. Adult female olfaction assay

The adult female olfaction assay was completed using protocols described previously (Wulff et al., 2024), and by using a disposable static olfactometer adapted from (Martin et al., 2020) (SM7). Tested flies were 10-day-old gravid females of the LA07 *wt* and *LcNPF^-/-^* colonies. Ten flies, either of the *wt* or *LcNPF^-/-^* colony were released for each trial until complete n = 25 replicates per genotype (total = 250 flies per genotype). Flies were lured with 1 g of fresh and 1 g of rotten 73/27% beef to assess olfactory choice. Beef type and conditions were according to (Wulff et al., 2024).

#### 2.5.3. Oviposition substrate preference

To test oviposition substrate preference, a single *wt* female was placed in a rearing bottle overnight with four substrates representing all combinations of beef type (fresh or rotten) and lean-fat ratio (73/27% or 93/7%). Each of the four types of beef (5 g) was placed in 35 mm petri dishes and offered at RT (SM8). The assay was also conducted for *wt* and *LcNPF^-/-^*females using only two choices (fresh and rotten 73/27% beef). A total of n = 20-25 replicates were completed for each genotype.

#### 2.5.4. Male mating competition

Competition experiments were performed by mixing ten *wt* males and ten *LcNPF*^-/-^ males with ten *wt* virgin females. Approximately 16 h later, the flies were anesthetized using carbon dioxide and the females were collected and placed individually in glass scintillation vials with cotton bungs. For egg collection, the individual females were transferred to egg laying chambers (5.5 oz plastic disposable cup that had a hole cut at the bottom large enough to fit over a 35 mm petri dish). A small quantity of fresh 93/7% fresh beef had been placed in the petri dish. The females were allowed to lay eggs overnight and developing embryos were examined for the presence or absence of green fluorescence to assess male parentage.

#### 2.5.5 Statistical Analysis

Fly fertility and oviposition substrate preference were evaluated using Fisher’s exact test, with the null hypothesis (H_₀_) stating no association between the two categorical variables. Egg hatching rate and larval survival were analyzed via ANOVA, while differences among groups for larval weight gain, locomotion, and pupation were assessed using the Mann-Whitney test. For the adult female olfaction assay, we applied the Kruskal-Wallis test, and in male mating competitiveness assays, the mating competitiveness index (MCI) was calculated as the number of females mated to mutant (fluorescent) males divided by the total number of mated females. All data were analyzed in SAS (Version 9.4, Cary, NC). Counts were pooled across replicates. Exact and asymptotic confidence limits were calculated for the pooled MCI as well as separately for each replicate. The overall MCI was tested against the null-hypothesized value of 0.5 using a two-tailed z-test for a proportion. Results were considered to be statistically significant when *P* < 0.05.

## 3. Results and Discussion

### 3.1. Generation of a loss-function mutation in the *LcNPF* gene

To facilitate the easy distinction of live heterozygous and homozygous mutant larvae we used CRISPR/Cas9 to insert a constitutively expressed ZsGreen fluorescent protein gene into *LcNPF* as done previously for the *L. cuprina nbl* (Davis et al., 2018) and *Lctra* genes (Williamson et al., 2021) (Fig. 1). *Lucilia c. cuprina* precellular embryos were injected with Cas9/gRNA complex and plasmid DNA. The plasmid contained the *Lchsp83-ZsGreen* gene (Concha et al., 2011) flanked by approximately 1 kb homology arms that match the *LcNPF* gene sequence (Fig. 1). The Cas9 cut site is at the beginning of the CDS that encodes the signal peptide of the preproprotein (Fig 1C). Consequently, a knock-in mutation is expected to be a null mutation. Homozygotes identified by bright whole-body fluorescence (SM4), were viable and fertile, and further confirmed by PCR with one primer in the flanking DNA and one primer in the transgene (SM5). The homozygous *LcNPF* mutant line was then used for studies on gene expression, behavior and development. For most experiments, *LcNPF*^-/-^ homozygotes were compared to the parental wild type LA07 strain. For gene expression analysis, *LcNPF*^-/-^ were compared to *LcNPF*^+/-^ heterozygotes (instead of using the *wt* strain), to control any potential impact of transgene integration and expression on genome expression.

### 3.2. RNA-Seq overview, gene expression and differential expression

To gain an insight into the potential impact of loss of a functional *LcNPF* gene, we first performed whole-body RNA-Seq analyses. For comparison, sibling heterozygous *LcNPF*^+/-^ larvae were also collected on days 3 (D3) and 5 (D5) after egg hatching. RNA-Seq output including reads per library, total number of sequences mapped and subtotals per type of RNA are provided in Data S1A-B. Gene expression analysis for larvae collected at D3 and D5, are provided in Data S2A-L and S3A-L, respectively. The first two tabs of each file compile all the libraries for each group, either heterozygous or homozygous, and include only transcripts surpassing a 5 transcript per million (TPM) expression threshold. The remaining tabs include the results for each individual library within each group. In addition, alignment of reads to the *LcNPF* gene locus confirmed disruption of gene expression in homozygotes (Fig. S1A-G).

Differential gene expression (DGE), associated with the functional disruption of the *LcNPF* gene was analyzed, and results are provided in Data S4. Differentially expressed transcripts in larvae collected on D3, exhibiting upregulation (Log_₂_FC > 1) or downregulation (Log_₂_FC < −1) were placed in Data S4A. Further, those transcripts found differentially expressed in larvae collected on D5 (identified using the same criteria as above), are shown in Data S4B and S4C. To enhance clarity, these transcripts were divided into two separate tabs, where B includes the downregulated, and C the upregulated transcripts. Down and upregulated transcripts were those showing a decreased or increased expression, respectively, in HOM NPF null mutated larvae in comparison to HET larvae, at any developmental time.

The DGE analysis produced 156 differentially expressed transcripts in larvae collected on D3 (a mix of late L2 and early L3 larvae). In contrast, 2,892 transcripts were differentially expressed in those larvae collected on D5, including only larvae from the late L3 larval stage. These findings may suggest a paramount regulatory role of *LcNPF* during the late larval developmental time. This observation was also supported by a Principal Component Analysis (PCA, Fig. S2A-B), where HET and HOM samples were completely separated only for larvae collected on D5. Interestingly, when analyzing transcript expressions across developmental stages of *L. cuprina wt* larvae, we previously observed that *LcNPF* transcript levels were significantly higher in early larval stages than in later ones (Wulff et al., 2025). The discrepancy with our findings may reflect a temporal offset between transcript abundance and peptide activity. In insects, numerous neuropeptides are known to be stored in vesicles within neurosecretory cells, and their release and action may occur independently of transcript expression timing (Bulgari et al., 2014; Gilbert and Rybczynski, 2008).

Among the most affected physiological processes after functional disruption of *LcNPF* in larvae collected on D5, included immune response (receptors and effectors), metabolism (carbohydrate, lipid and protein), cuticle formation and sensory and detoxification (Fig. 2). The functions attributed to each transcript were defined according to literature screening (added to the reference section) and NCBI annotations. These mentioned processes are discussed in more detail within each respective section below.

**Figure 2.**
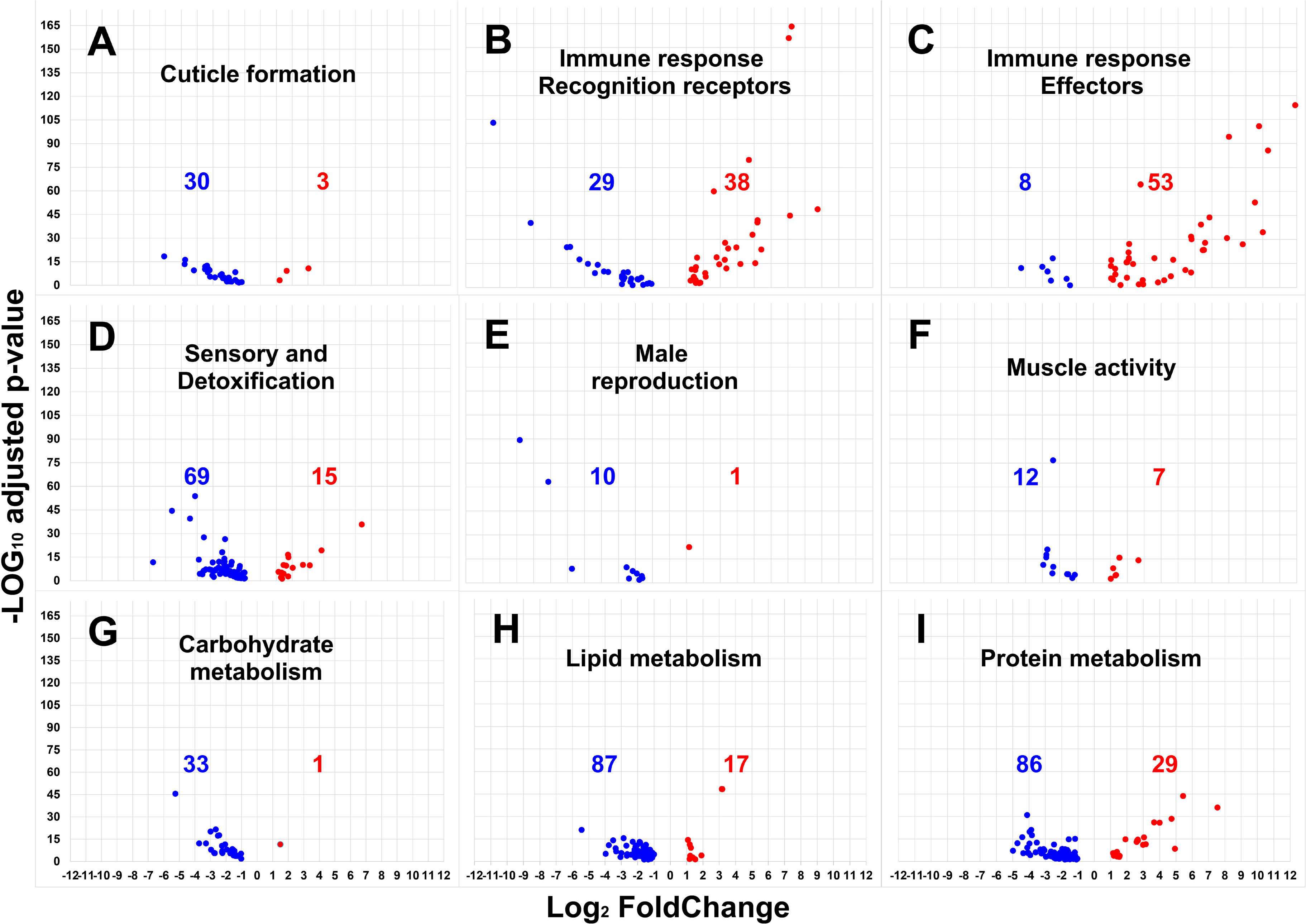
Combined volcano plot showing down- and upregulated transcripts after functional disruption of the *LcNPF* gene in NPF null mutated larvae. The most impacted biological processes were determined based on the magnitude of differential gene expression and the number of transcripts per gene family (Data S4B and S4C). **A**. Cuticle formation. **B**. Immune response linked to pathogen recognition, *e.g*. peptidoglycan. **C**. Immune response effectors, *e.g*. antimicrobial peptides (AMPs). **D**. Sensory and detoxification, *e.g*. cytochrome P450s and odorant-binding proteins among others. **E**. Male reproduction. **F**. Muscle formation and contraction. **G**. Carbohydrate metabolism, *e.g*. transcripts involved in glycolysis and gluconeogenesis. **H**. Lipid metabolism, *e.g*. sequences associated with lipolysis, lipogenesis and lipid transport. **I**. Protein metabolism, *e.g*. peptidases, proteases and protein storage. The y-axis shows statistically significant as −Log_10_ *P*-adjusted value and the x-axis, down- and upregulated transcripts in homozygous compared to heterozygous larvae collected on day 5 (five libraries per group), presented in Log_2_ fold-change. Down- and upregulated transcripts are in blue and red respectively. Numbers represent the biased transcripts within each group.

### 3.3. *Loss of NPF* doesn’t affect male mating fitness and female fertility, but it does impair the egg hatching rate

We next examined male mating competitiveness as *npf* is essential for normal male courtship behavior in *Drosophila* (Liu et al., 2019) and ten transcripts associated with the function of testes and male accessory glands were downregulated in D5 *LcNPF^-/-^* (Fig. 2 and Data S4B). Competition experiments were performed by mixing ten 10 *wt* males and ten *LcNPF^-/-^* males with ten *wt* females. After 16 h, females were collected and placed individually in egg laying chambers with ground meat. The male parentage could be determined by the presence or absence of green fluorescence in developing embryos. The overall MCI obtained from 11 replicate experiments was 0.49 (Table S1), which did not differ significantly from the expected value of 0.5 for fully competitive males. Further, the *LcNPF^-/-^* males showed normal fertility (Fig. S3). Thus, these results indicate that, unlike in *Drosophila*, NPF is not essential for normal male mating behavior in *L. cuprina*.

We also investigated the hatching rate of *LcNPF^-/-^* eggs under both normal and desiccation conditions. Under normal conditions, the average hatching rate was 81% and 57% for *wt* and *LcNPF^-/-^* eggs respectively (Fig. S4). Using a 5 min desiccation treatment immediately after egg collection, the hatching rate was 81% and 62% for *wt* and *LcNPF^-/-^*eggs respectively (Fig. S4). Our gene expression results showed numerous genes involved in cuticle formation that were downregulated in *LcNPF^-/-^*larvae collected on D5 (Fig. 2), such as the *vitelline membrane protein Vm26Ab-like* gene (see Data S4B). In *D. melanogaster*, *Vm26Ab* is involved in vitelline membrane biogenesis and is essential for female fertility (Wu et al., 2010). However, our findings indicated that female fertility was normal (Fig. S3), and desiccation was unlikely to account for the reduced hatching rate observed in *LcNPF^-/-^* eggs.

In *Schistocerca gregaria* Forsskål, 1775 (Orthoptera: Acrididae), the disruption of the NPF signaling also affected egg hatching (Van Wielendaele et al., 2013). It was suggested that the low hatching egg may be linked to the quantity and quality of sperm transferred by null mutated NPF males (Van Wielendaele et al., 2013). However, fertility of *LcNPF^-/-^*males was comparable to wild type in our experiments, and consequently, further studies are necessary to determine the cause associated with the low hatching rate of *LcNPF^-/-^* eggs.

### 3.4. *LcNPF* participates in larval development and weight gain

Larval weight gain was assessed at two larval feeding stages, early- (2-day-old) and late-L2 (3-day-old), as well as for the non-feeding stage, the wandering L3 (6-day-old). In addition, two types of ground beef were used to rear the larvae as follows, 73/27% and 93/7% lean/fat. Both *wt* and *LcNPF^-/-^* larvae showed greater weight gain using 93/7% beef (high protein diet) (Fig. 3). On both diets and at all stages the weight of *wt* larvae was greater than *LcNPF^-/-^*larvae. The difference in weight was most pronounced in late L2 on 93/7% beef (1.6-fold). The gene expression results supported these findings, as many transcripts associated with carbohydrate, lipid and protein metabolism were downregulated in *LcNPF^-/-^* larvae collected on D5 (Fig. 2G-I and Data S4B). In addition, the downregulated genes included enzymes associated with food-protein digestion by *L. cuprina* larvae, such as trypsin and chymotrypsin (Hobson RP, 1932). Interesting, the same enzymes had been previously identified as upregulated in the second instar of *L. cuprina* compared to the L1 and L3 stages (Wulff et al., 2025) which may explain the large weight difference observed between the *wt* and *LcNPF^-/-^* genotypes at the late-L2 larval stage (Fig. 3B). Moreover, the difference in average weight between *wt* and *LcNPF^-/-^* larvae was higher at all stages when using 93/7% beef (Figs. 3A-C vs 3D-F), highlighting the key role of the enzymes to digest food with a high protein content.

**Figure 3.**
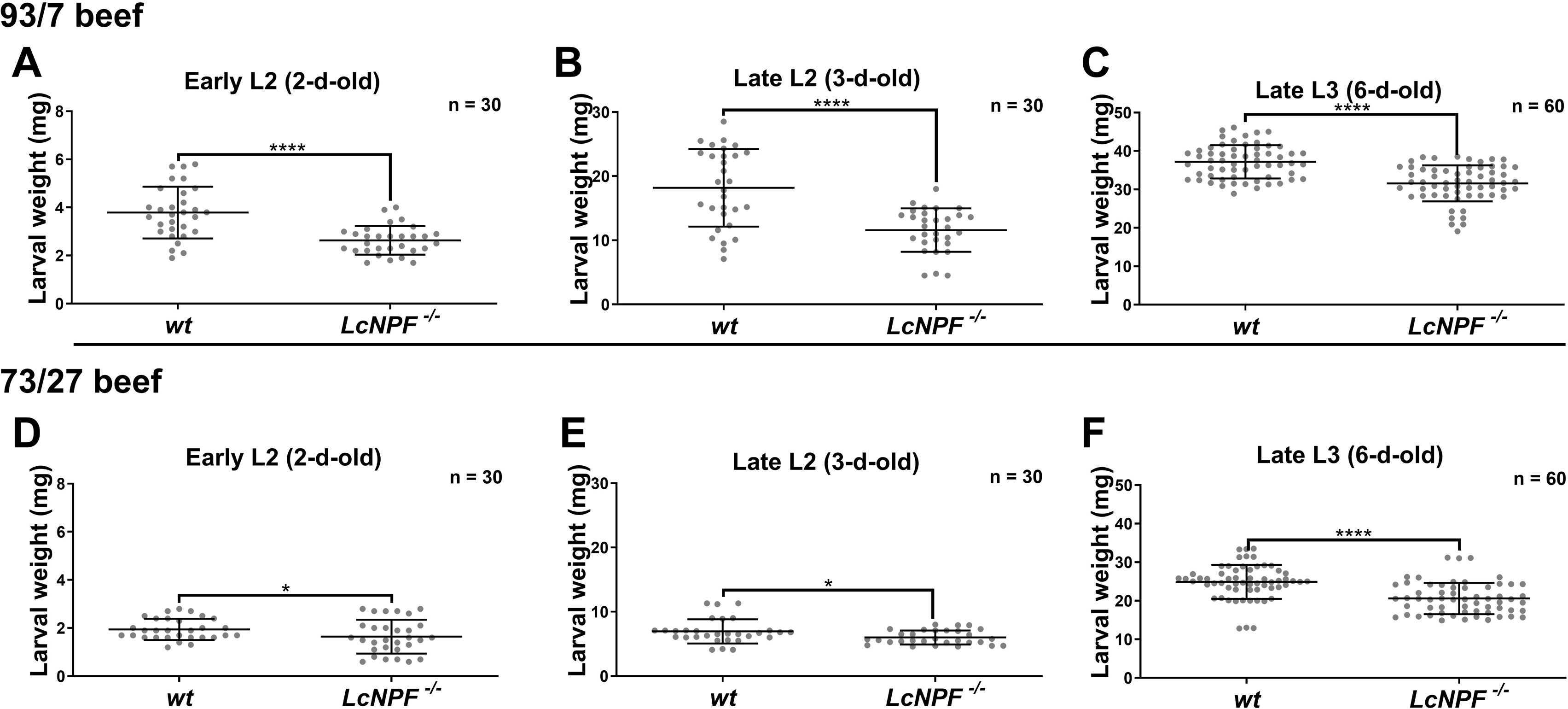
Weight of wild type (*wt*) and *LcNPF* mutant (LcNPF^-/-^) feeding and non-feeding wandering larvae. Feeding larvae at the early (2 days) (**A**, **D**) or late (3 days) (**B**, **E**) second-instar stage (L2) were removed from diet and weighed. Wandering third-instar larvae (L3) were also collected and weighed (**C**, **F**). The diets used were 93/7% lean-fat (**A**-**C**) and 73/27% (**D**-**F**) ground fresh beef. Results were plotted as mean ± standard deviation (SD). Statistically significant differences were tested using Mann-Whitney test with a sample size n = 30 for both L2 stages and n = 60 for L3. One, two, three and four asterisks correspond to *P*-values of < 0.05, < 0.01, < 0.001, and < 0.0001 respectively.

We next explored the time to pupariation of the wandering L3 stage and the adult emergence of *L. cuprina.* Both the pupariation time (Fig. S5A) and time to adult emergence (Fig. S5B) were statistically different between the *wt* and *LcNPF^-/-^* genotypes. In *D. melanogaster* NPF signaling regulates the time to pupariation by controlling the prothoracic gland activity and consequently, the ecdysone biosynthesis, and its interruption was associated with a delay in completing this process (Kannangara et al., 2020). In addition, NPF has been demonstrated to influence the juvenile hormone (JH) and 20-hydroxyecdysone (20E) biosynthesis pathways in other insects (Kannangara et al. 2020; Yu et al. 2023). However, *LcNPF^-/-^*individuals showed an acceleration of the time to pupariation and adult emergence (Fig. S5A-B). Furthermore, genes associated with JH biosynthesis and the ecdysone conversion into its active form (20E), such as *farnesol dehydrogenase-like* and *ecdysone 20-monooxygenase* respectively, were downregulated in *LcNPF^-/-^* larvae collected on D5 (Data S4B). The same analysis also revealed the upregulation of genes implicated in ecdysone biosynthesis and ecdysone-induced transcription factors (Erkelenz et al., 2021; Stone and Thummel, 1993), such as *protein ecdysoneless* and *ecdysone-induced protein 78C* respectively, in NPF null mutated individuals (Data S4C). Our findings indicate that NPF may function as a modulator of this process rather than serving as a direct enhancer or repressor, a similar role observed in association with immune response in *D. melanogaster* (Deng and Chiu, 2022).

### 3.5. *LcNPF* disruption decreased larval crawling speed

Locomotion performance of the wandering stage was evaluated (Post and Paululat, 2018), and there was 24% decrease in larval crawling speed in *LcNPF^-/-^*in comparison to *wt* larvae (Fig. 4). This result is consistent with the gene expression results that showed numerous genes associated with muscle function downregulated in the *LcNPF^-/-^*larvae collected on D5, such as myosin, paramyosin and troponin (Fig. 2F, Data S4B). In *D. melanogaster*, NPF receptor 1 (NPFR1) has been identified in motor neurons innervating the abdominal muscles (Wu et al., 2003), and NPF has been shown to regulate locomotor activity in both larval and adult stages (Hermann et al., 2012; Wu et al., 2003), but no locomotory muscle defects affecting movement have been reported (Wu et al., 2003). However, the myotropic activity of NPF has been confirmed in the musculature of gonads and heart (Liu et al., 2021), the latter, in adult males of the blowfly *Pr. terraenovae*. These findings suggest that NPF may play a role in regulating muscle contraction across different tissues and developmental stages, a hypothesis that will be further investigated in future studies.

**Figure 4.**
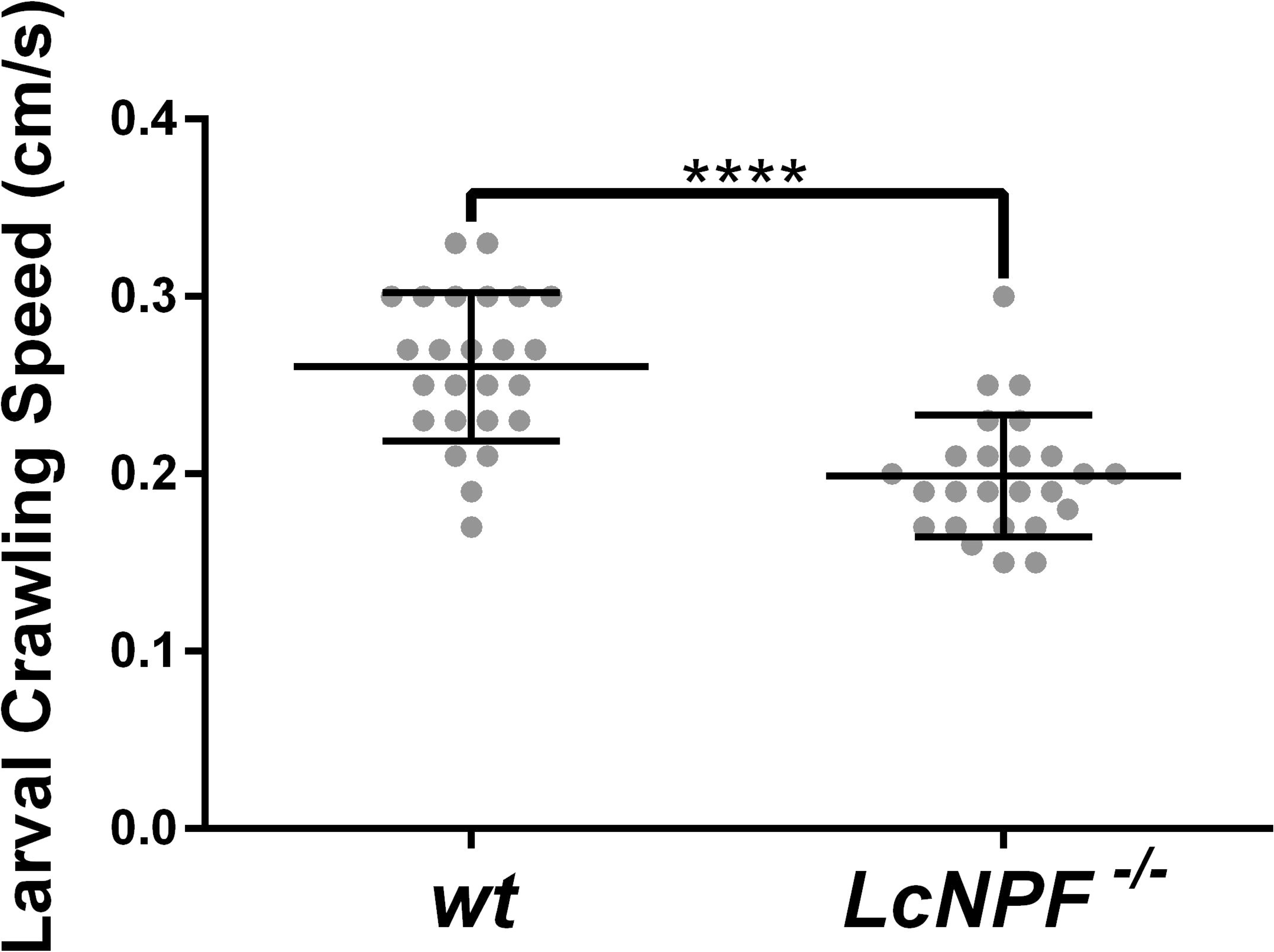
Larval crawling speed of wild type (*wt*) and *LcNPF* homozygous (*LcNPF^-/-^)* wandering L3 larvae. Results were plotted as mean ± standard deviation (SD). Statistically significant differences were tested using unpaired t-test test with a sample size n = 25 and a *P* < 0.0001.

### 3.6. *LcNPF* affects larval survival on rotting beef

The *wt L. c. cuprina* subspecies used in this study is necrophagous and not associated with myiasis (Kapoor et al., 2025a). We previously found that *wt* LA07 larvae and adult females are strongly attracted to rotting beef (Cunha et al., 2023; Wulff et al., 2024). We next investigated if *LcNPF* played a role in larval survival on fresh and rotting beef. Larval survival at L2-L3 stages was evaluated using fresh and rotten 93/7% and 73/27% lean-fat ground beef (Table S2 and S3). The rotten beef showed a detrimental effect to both *wt* and *LcNPF^-/-^* larvae, as evidenced by reductions in the number of surviving larvae, the proportion that progressed to the pupal stage, and the pupal emergence rate (Fig. 5A-D). However, the negative impact was greatest for *LcNPF^-/-^* larvae (Fig. 5B-D), which completely failed to reach the pupal and adult stages when reared using the rotten fatty 73/27% lean-fat beef (Fig. 5D). Among the beef types tested, the fatty beef exhibited a more pronounced negative impact on both genotypes (Fig. 5C-D). This may be attributed to its nutritional inadequacy which appears to be less suitable for the development of *L. cuprina* larvae, as demonstrated in Fig. 3.

**Figure 5.**
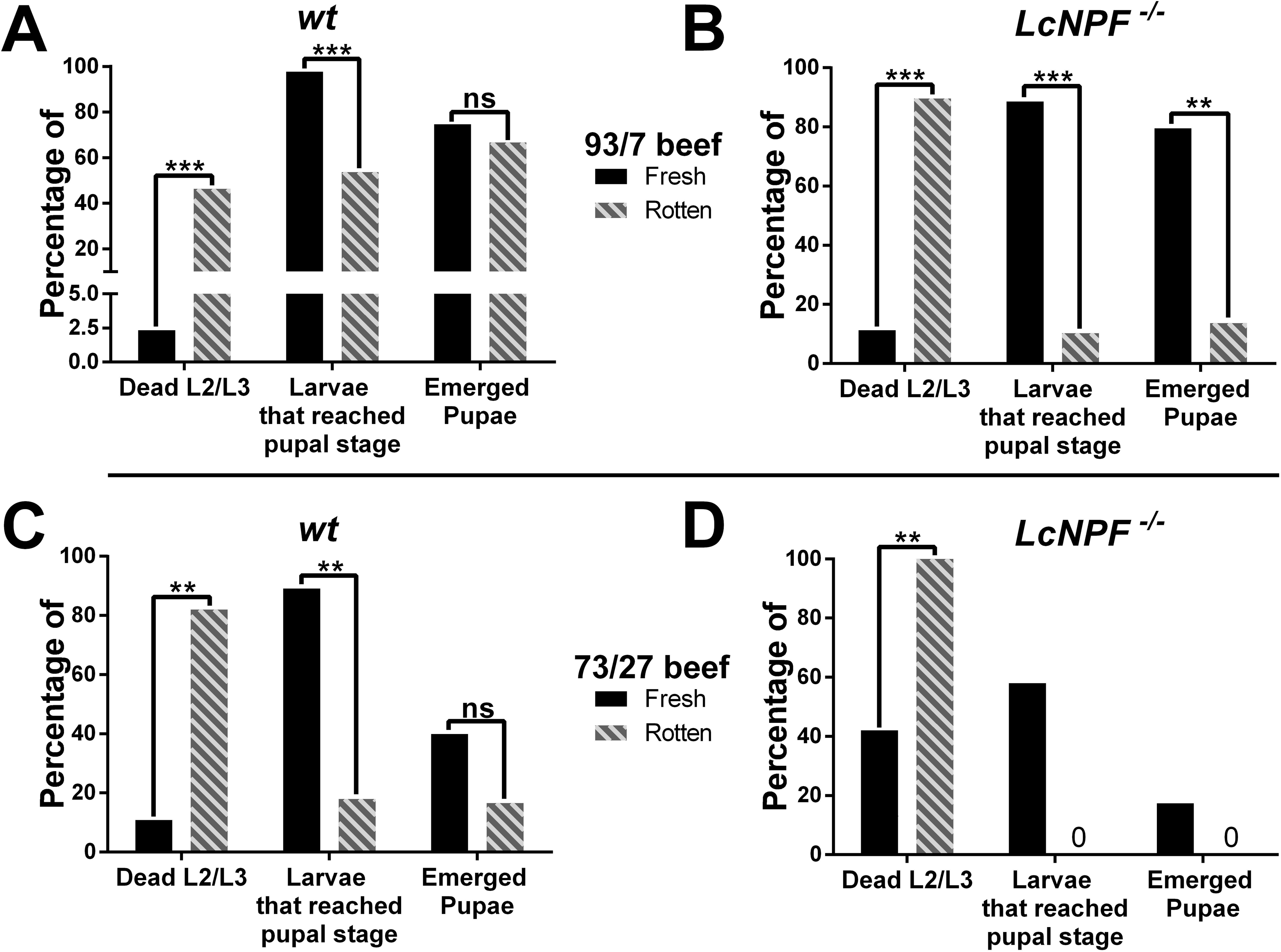
Larval survival when reared on fresh and rotting lean and fatty ground beef. The percentage of L2 and L3 that died, the proportion that progressed to the pupal stage, and the pupal emergence rate are shown for *wt* (**A**, **C**) and *LcNPF^-/-^* homozygotes (**B**, **D**) on fresh and rotten 93/7% (**A**, **B**) and 73/27% (**C**, **D**) ground beef. Results were plotted as percentages and for more details refer to Table S1. Statistically significant differences were tested using Multiple t-test test with a sample size n = 3 replicates per condition. Replicates were pools and the number of larvae per pool is provided in Table S2. Abbreviation ns corresponds to non-statistically significant, and two or three asterisks to *P*-values of *P* < 0.01 and *P* < 0.001, respectively.

In *D. melanogaster*, NPF null mutant flies showed high mortality following exposure to Gram-negative bacteria (Deng and Chiu, 2022). This mortality was attributed to autoimmune effects since many genes associated with immune response regulation and effectors, such as *eiger* and antimicrobial peptides respectively, were upregulated during the assay (Deng and Chiu, 2022). Consequently, NPF appears to play a role in modulating an adequate immune response. These findings are consistent with our gene expression results, which revealed significant upregulation of numerous genes associated with immune response in larvae collected on D5 (Data S4C), like those included in the Imd-pathway, such as *relish* (De Gregorio, 2002). However, the same analysis also revealed downregulation of other immune genes (Data S4B), such as spätzle, belonging to the Toll-pathway (De Gregorio, 2002). Toll and Imd pathways primarily drive the immune response to Gram-positive and -negative bacteria respectively (De Gregorio, 2002), and we plan to complete future experiments using different types of bacteria to evaluate the role of NPF in the immune response of *L. c. cuprina*.

Additionally, a consequence of a high protein diet is the production of nitrogenous waste, which is a major problem for the mass rearing of *Ch. hominivorax* larvae (Chen et al., 2014). The gene expression results revealed that the *ammonium transporter Rhesus type B* (*RHBG*) was significantly downregulated in NPF null mutated larvae collected on D5 (Data S4B). This transporter is thought to be important for nitrogenous waste excretion (Durant and Donini, 2024) and was recently found to be overexpressed in the gut of *L. c. cuprina* larvae (Wulff et al., 2025). To investigate the functional implications of this downregulation, we supplemented fresh meat with two nitrogen sources, ammonium hydroxide or urea (Table S3). However, neither treatment produced a deleterious effect on larval development (Fig. S6A-B). These findings suggest that nitrogen compounds excretion in *L. cuprina* may involve a redundant genetic system, as observed in other dipteran species such as those of the Culicidae family, including genes like alanine aminotransferases and urate oxidase, involved in nitrogen homeostasis (Durant and Donini, 2024).

### 3.7. *LcNPF* disruption alters female olfaction, but not larval feeding preference behavior

We evaluated the consequence of *LcNPF* disruption on female olfactory and oviposition preference behavior and larval feeding preference behavior. Previous evidence demonstrated that NPF signaling is involved in host detection and female oviposition preference in *Aedes aegypti* Linnaeus, 1762 (Diptera: Culicidae) and *Spodoptera frugiperda* Smith, 1797 (Lepidoptera: Noctuidae) (Dou et al., 2024; Xie et al., 2025), and larval foraging in the latter (Xie et al., 2025). *Lucilia c. cuprina wt* females were more attracted to rotten beef when fresh and rotten beef was offered to them (Wulff et al., 2024). However, *LcNPF^-/-^* mutant females did not show this characteristic olfactory response being equally attracted to fresh and rotten 73/27% beef (Fig. 6 and Table S4).

**Figure 6.**
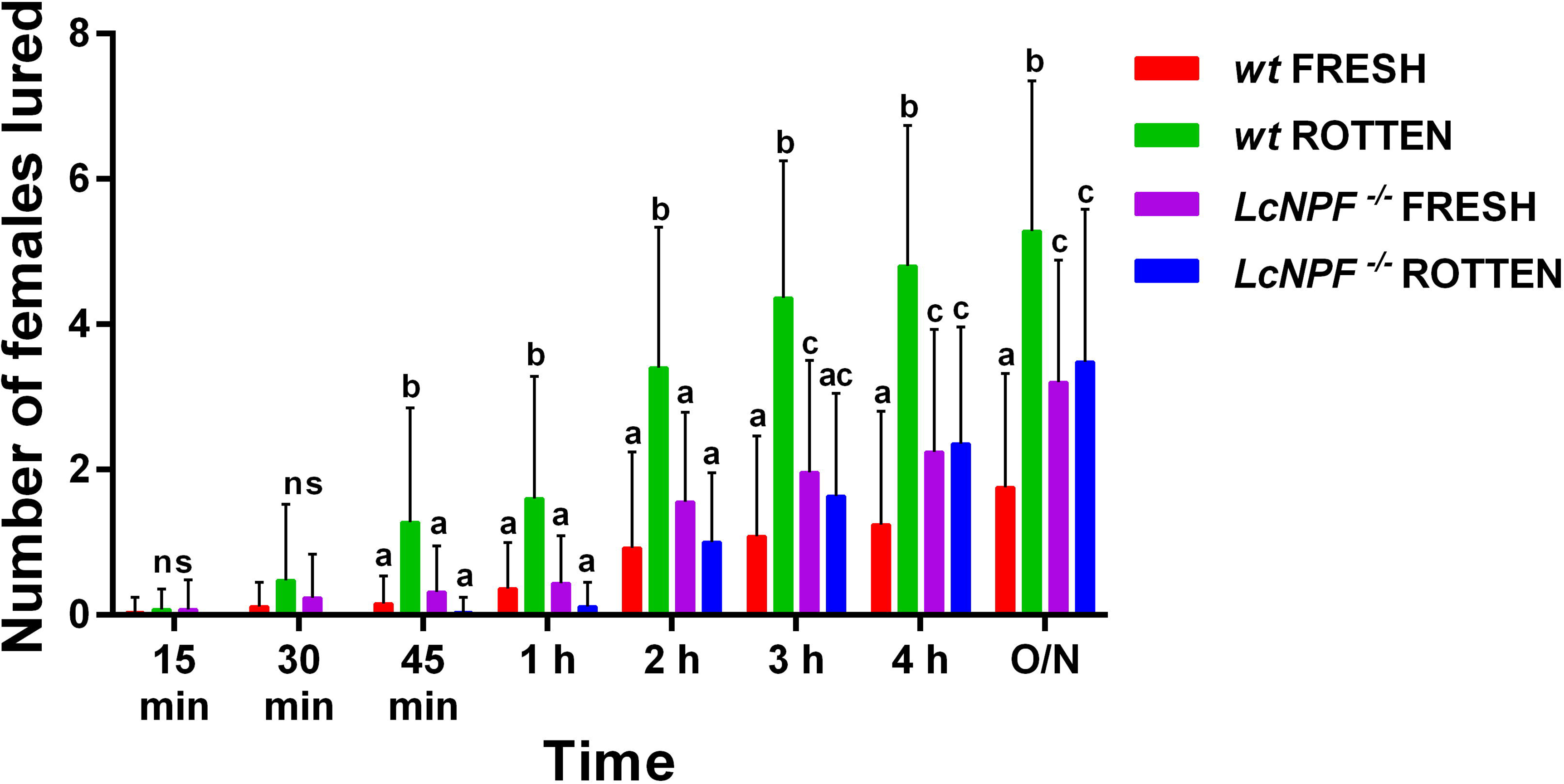
*LcNPF* disruption alters adult’s female olfaction. The number of *wt* and *LcNPF* mutant (*LcNPF^-/-^*) females that were attracted to fresh or rotten beef in a static olfactometer over time are shown. Statistically significant differences were tested using repeated-measures one-way ANOVA with a sample size n = 25 replicates per genotype, including 10 females per replicate. Abbreviation ns corresponds to non-statistically significant, and different lowercase letters mean that the compared columns were statistically significantly different. For more details refers to Table S4.

Additionally, we assessed the oviposition preferences of *L. cuprina* females across various substrates (Fig. 7A and Table S5). In this assay, *wt* females exhibited a preference for rotten 73/27% beef, whereas NPF null mutated females showed a marked preference for fresh 73/27% beef (Fig. 7B and Table S6). Notably, while *wt L. cuprina* females exhibit a preference for rotten beef as an oviposition substrate, larval survival on this medium was markedly low (Fig. 5). Previous studies have suggested that Calliphoridae larvae must be present in sufficient densities to secrete adequate levels of secretions, including ammonia and antimicrobial peptides, to suppress harmful bacterial populations (Yoon et al., 2022). In our larval survival assay, only 200-300 larvae were placed per sample on either fresh or 5-day-rotten beef (Table S2-S3), the latter carrying a substantial bacterial load. This limited larval density may account for the reduced survival observed on the rotting substrate.

**Figure 7.**
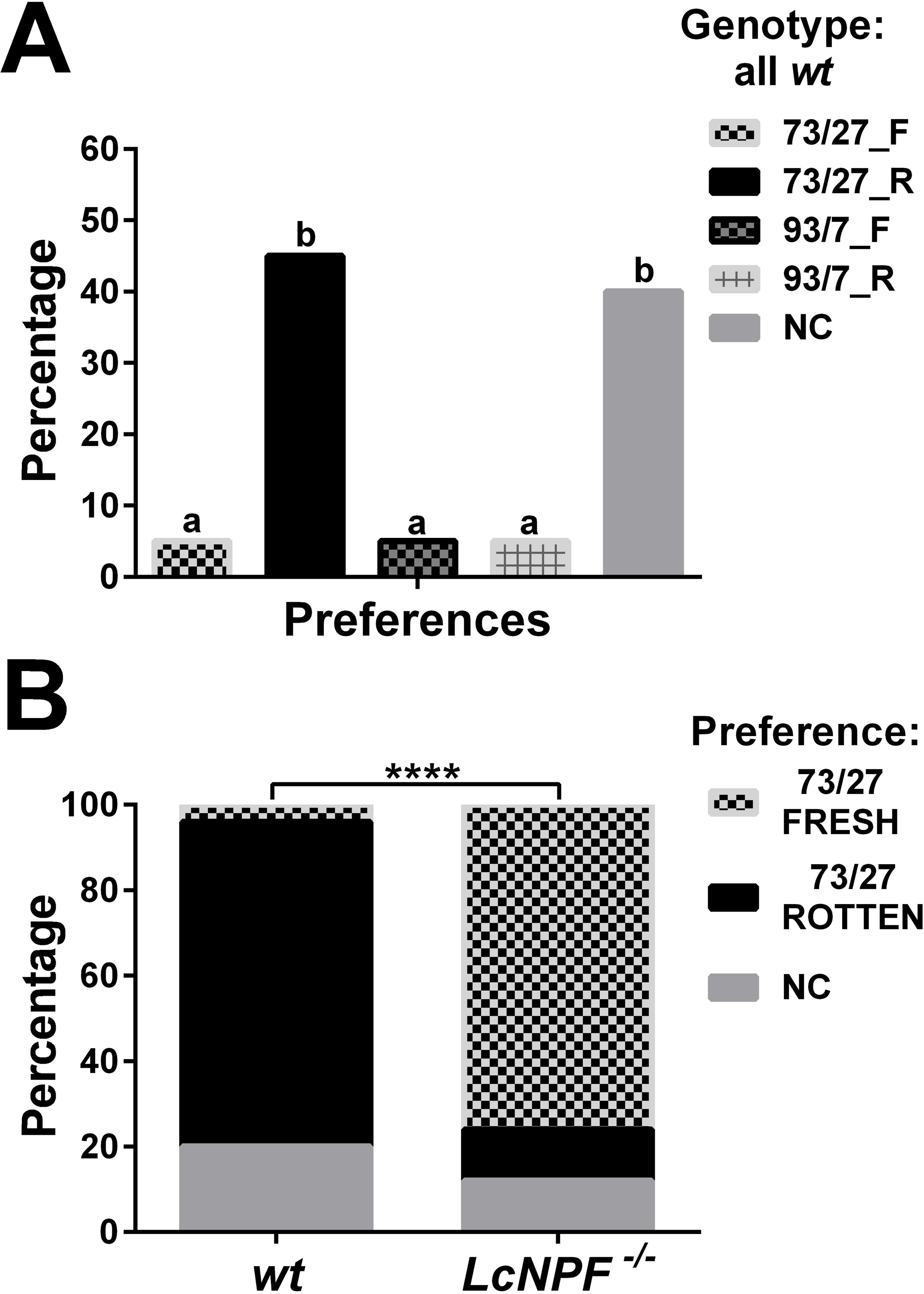
*Lucilia cuprina* female’s oviposition preference across different beef substrates. *Lucilia cuprina* wild-type (*wt*) females exhibited a preference for fatty (73/27% lean-fat) rotten beef (**A, B**), whereas neuropeptide F (*LcNPF^-/-^*) null mutated females showed a marked preference for fresh beef, when tested using fresh and rotten fatty beef (**B**). Statistically significant differences were tested using Mann-Whitney test with a sample size n = 20 replicates of *wt* flies (A) or n = 25 replicates per genotype (**B**). Different lowercase letters mean that the compared columns were statistically significant different, and four asterisks correspond to *P*-values of *P* < 0.0001. For more details, refer to Table S5 and S6.

In contrast to adult females, *LcNPF^-/-^* larvae (mix of late L2/early L3) showed no differences in feeding preference behavior when offered fresh or rotten beef at 25 ± 1 or 33 ± 1°C compared to the *wt* larvae (Fig. S7A-B). Interestingly, transcript levels of both NPF and its receptor were found to be very low in the adult female antennae and the anterior region of larvae (Wulff et al., 2025, 2024). In this regard, studies in *D. melanogaster* have demonstrated that NPF modulates peripheral olfactory responses in both adults and larvae via a central peptidergic circuit located in the brain (Lee et al., 2017; Pu et al., 2018). These findings suggest that NPF may have a comparable physiological function in *L. cuprina* but possibly restricted to adult flies. In this regard, in *D. melanogaster* NPF has been shown to modulate the spatial localization of odorant receptor 22a within sensory neurons (Lee et al., 2017). Notably, our previous findings revealed that members of the odorant receptor gene family are minimally expressed during the larval stage of *L. cuprina* (Wulff et al., 2025) yet are prominently expressed in the adult antennae (Wulff et al., 2024). Consequently, NPF modulation of olfactory responses in *L. cuprina* may be through the regulation of odorant signaling and be more or only relevant for adult flies.

A parasitic lifestyle appears to have evolved independently multiple times in Calliphoridae (Cardoso et al., 2025; Nasser et al., 2021). The *wt L.c. cuprina* subspecies from the New World used in this study is necrophagous whereas the *L. c. dorsalis* subspecies causes flystrike in sheep in Australia and New Zealand and is a major economic pest (Kapoor et al., 2025b). Disruption of the *LcNPF* gene caused a significant shift in oviposition preference from rotting meat to no choice between fresh and rotten (Fig. 6). Changes in oviposition preference have been hypothesized to play a crucial role in the evolution of parasitism in blow flies (Cardoso et al., 2025). Consequently, it would be of interest to determine if *LcNPF* plays a role in the oviposition preference of *L. c. dorsalis* females and to compare *LcNPF* gene expression in the two *L. cuprina* subspecies.

## Supporting information

type of RNA are provided in

D3 and D5, are provided in

and S3A-L, respectively

are provided in Data S4. Differentially expressed

MCI obtained from 11 replicate experiments was 0.49 (Table S1)

in the supplementary method (SM) section SM1A-C

straight-line crawling (Video S1) were

## Data availability

All relevant data are in the manuscript. Raw sequences were uploaded to the NCBI database under Bioproject PRJNA1372265.

## Author contributions

**Juan P. Wulff**: Conceptualization, Methodology, Investigation, Visualization, Writing-Original draft preparation. **Esther J. Belikoff**: Methodology, Investigation. **Vanesa A.S. Cunha**: Methodology, Investigation. **Pedro Mariano-Martins**: Methodology, Investigation. **Maxwell J. Scott**: Conceptualization, Writing-Original draft preparation, Visualization, Supervision, Project administration, Funding acquisition.

## Acknowledgements

The authors thank Emily Griffith for statistical analysis of the male mating competition data.

## Supplementary data legends

**Data S1**. RNA-Seq output, including in tab A reads per library, total number of sequences trimmed and mapped and subtotals per type of RNA and cytosine-guanine % in tab B.

**Data S2**. Gene expression analysis of L2 collected 3 days after egg hatching. The first two tabs A-B compile all the libraries for each group (heterozygous or homozygous), including only transcripts surpassing a 5 transcripts per million (TPM) expression.

**Data S3**. Gene expression analysis of L3 collected 5 days after egg hatching. The first two tabs A-B compile all the libraries for each group (heterozygous or homozygous), including only transcripts surpassing a 5 transcripts per million (TPM) expression.

**Data S4**. Differential gene expression analysis, where tab A corresponds to L2 and B-C to L3 larvae. Differentially expressed (DE) transcripts for L2 larvae, exhibiting a Log_₂_ fold-change (Log_2_FC) greater than 1 or less than −1 and a TPM value ≥ 5, were placed in tab B and DE transcripts for L3 larvae selected using the same criteria are shown in tabs B and C. Tab B contains downregulated transcripts (Log_₂_FC < −1), while tab C contains upregulated transcripts (Log_₂_FC > 1).

**Video S1**. Single L3 wandering larvae exhibiting a straight-line crawling path as used for the larval locomotion assay.

## References

Amaral Faria, G., 2019. Feeding habit in Calliphoridae (Insecta, Diptera, Oestroidea): bionomy and molecular basis. Universidade Estadual de Campinas, Campinas. 10.47749/T/UNICAMP.2019.1126965

Angioy, A.M., Muroni, P., Barbarossa, I.T., McCormick, J., Nichols, R., 2007. Evidence dromyosuppressin acts at posterior and anterior pacemakers to decrease the fast and the slow cardiac activity in the blowfly *Protophormia terraenovae*. Peptides (N.Y.) 28, 585–593. 10.1016/j.peptides.2006.10.015

Ashworth, J.R., Wall, R., 1994. Responses of the sheep blowflies *Lucilia sericata* and *L. cuprina* to odour and the development of semiochemical baits. Med Vet Entomol 8, 303–309. 10.1111/j.1365-2915.1994.tb00093.x

Benbow, M.E., Tomberlin, J.K., Tarone, A.M., 2015. Carrion Ecology, Evolution, and Their Applications, Carrion ecology, evolution, and their applications. CRC Press. 10.1201/b18819

Bulgari, D., Zhou, C., Hewes, R.S., Deitcher, D.L., Levitan, E.S., 2014. Vesicle capture, not delivery, scales up neuropeptide storage in neuroendocrine terminals. Proceedings of the National Academy of Sciences 111, 3597–3601. 10.1073/pnas.1322170111

Byrd, J.H., Tomberlin, J.K., 2019. Forensic entomology: the utility of arthropods in legal investigations. CRC press.

Cardoso, G.A., Mariano-Martins, P., Faria, G.A., Karunaratne, I., Thyssen, P.J., Torres, T.T., 2025. Divergent Genetic Pathways Underlying Convergent Parasitic Behaviours in Blowflies. Mol Ecol 34. 10.1111/mec.17785

Casu, R.E., Pearson, R.D., Jarmey, J.M., Cadogan, L.C., Riding, G.A., Tellam, R.L., 1994. Excretory/secretory chymotrypsin from *Lucilia cuprina*: purification, enzymatic specificity and amino acid sequence deduced from mRNA. Insect Mol Biol 3, 201–211. 10.1111/j.1365-2583.1994.tb00168.x

Cerstiaensa, A.N.J.A., Benfekihb, L., Zouitenc, H., Verhaerta, P., De Loofa, A., Schoofsa, L., 1999. Led-NPF-1 stimulates ovarian development in locusts. Peptides (N.Y.) 20, 39–44. 10.1016/S0196-9781(98)00152-1

Chaudhury, M.F., Skoda, S.R., Sagel, A., Welch, J.B., 2010. Volatiles Emitted From Eight Wound-Isolated Bacteria Differentially Attract Gravid Screwworms (Diptera: Calliphoridae) to Oviposit. J Med Entomol 47, 349–354. 10.1603/ME09235

Chen, H., Chaudhury, M.F., Sagel, A., Phillips, P.L., Skoda, S.R., 2014. Artificial diets used in mass production of the New World screwworm, *Cochliomyia hominivorax*. Journal of Applied Entomology. 10.1111/jen.12112

Chung, B.Y., Ro, J., Hutter, S.A., Miller, K.M., Guduguntla, L.S., Kondo, S., Pletcher, S.D., 2017. Drosophila Neuropeptide F Signaling Independently Regulates Feeding and Sleep-Wake Behavior. Cell Rep 19, 2441–2450. 10.1016/j.celrep.2017.05.085

Concha, C., Belikoff, E.J., Carey, B., Li, F., Schiemann, A.H., Scott, M.J., 2011. Efficient germ-line transformation of the economically important pest species *Lucilia cuprina* and *Lucilia sericata* (Diptera, Calliphoridae). Insect Biochem Mol Biol 41, 70–75. 10.1016/j.ibmb.2010.09.006

Concha, C., Palavesam, A., Guerrero, F.D., Sagel, A., Li, F., Osborne, J.A., Hernandez, Y., Pardo, T., Quintero, G., Vasquez, M., Keller, G.P., Phillips, P.L., Welch, J.B., McMillan, W.O., Skoda, S.R., Scott, M.J., 2016. A transgenic male-only strain of the New World screwworm for an improved control program using the sterile insect technique. BMC Biol 14. 10.1186/s12915-016-0296-8

Cui, H., Zhao, Z., 2020. Structure and function of neuropeptide F in insects. J Integr Agric 19, 1429– 1438. 10.1016/S2095-3119(19)62804-2

Cunha, V.A.S., Tandonnet, S., Ferreira, D.L., Rodrigues, A. V., Torres, T.T., 2023. Exploring Life History Choices: Using Temperature and Substrate Type as Interacting Factors for Blowfly Larval and Female Preferences. Journal of Visualized Experiments e65835. 10.3791/65835

Davis, R.J., Belikoff, E.J., Scholl, E.H., Li, F., Scott, M.J., 2018. no blokes Is Essential for Male Viability and X Chromosome Gene Expression in the Australian Sheep Blowfly. Current Biology 28, 1987–1992.e3. 10.1016/j.cub.2018.05.005

De Gregorio, E., 2002. The Toll and Imd pathways are the major regulators of the immune response in Drosophila. EMBO J 21, 2568–2579. 10.1093/emboj/21.11.2568

Deng, L., Chiu, I.M., 2022. A neuropeptide regulates immunity across species. Neuron 110, 1275–1277. 10.1016/j.neuron.2022.03.036

Doll, L.C., 2020. An investigation of genetic variability in *Lucilia cuprina* and *Musca domestica* utilizing phylogenetic and population genetic approaches.

Dou, X., Chen, K., Brown, M.R., Strand, M.R., 2024. Reciprocal interactions between neuropeptide F and RYamide regulate host attraction in the mosquito *Aedes aegypti*. Proceedings of the National Academy of Sciences 121. 10.1073/pnas.2408072121

Downer, K.E., Haselton, A.T., Nachman, R.J., Stoffolano, J.G., 2007. Insect satiety: Sulfakinin localization and the effect of drosulfakinin on protein and carbohydrate ingestion in the blow fly, *Phormia regina* (Diptera: Calliphoridae). J Insect Physiol 53, 106–112. 10.1016/j.jinsphys.2006.10.013

Durant, A.C., Donini, A., 2024. Ammonia transport in the excretory system of mosquito larvae (*Aedes aegypti*): Rh protein expression and the transcriptome of the rectum. Comp Biochem Physiol A Mol Integr Physiol 294, 111649. 10.1016/j.cbpa.2024.111649

Duve, H., Thorpe, A., 1994. Distribution and functional significance of Leu-callatostatins in the blowfly *Calliphora vomitoria*. Cell Tissue Res 276, 367–379. 10.1007/BF00306122

Duve, H., Thorpe, A., Lazarus, N.R., 1979. Isolation of material displaying insulin-like immunological biological activity from the brain of the blowfly *Calliphora vomitoria*. Biochemical Journal 184, 221–227. 10.1042/bj1840221

Erkelenz, S., Stanković, D., Mundorf, J., Bresser, T., Claudius, A.-K., Boehm, V., Gehring, N.H., Uhlirova, M., 2021. Ecd promotes U5 snRNP maturation and Prp8 stability. Nucleic Acids Res 49, 1688–1707. 10.1093/nar/gkaa1274

Fadda, M., Hasakiogullari, I., Temmerman, L., Beets, I., Zels, S., Schoofs, L., 2019. Regulation of Feeding and Metabolism by Neuropeptide F and Short Neuropeptide F in Invertebrates. Front Endocrinol (Lausanne) 10. 10.3389/fendo.2019.00064

Fenton, A., Wall, R., French, N.P., 1999. The effects of oviposition aggregation on the incidence of sheep blowfly strike. Vet Parasitol 83, 137–150. 10.1016/S0304-4017(99)00047-3

Fónagy, A., 2006. Insect Neuropeptides and their Potential Application for Pest Control. Acta Phytopathol Entomol Hung 41, 137–152. 10.1556/APhyt.41.2006.1-2.13

Forbes, S.L., Perrault, K.A., 2014. Decomposition Odour Profiling in the Air and Soil Surrounding Vertebrate Carrion. PLoS One 9, e95107. 10.1371/journal.pone.0095107

Gäde, G., Wilps, H., Kellner, R., 1990. Isolation and structure of a novel charged member of the red– pigment-concentrating hormone-adipokinetic hormone family of peptides isolated from the corpora cardiaca of the blowfly *Phormia terraenovae* (Diptera). Biochemical Journal 269, 309–313. 10.1042/bj2690309

Gilbert, L.I., Rybczynski, R., 2008. Prothoracicotropic Hormone, in: Encyclopedia of Entomology. Kluwer Academic Publishers, Dordrecht, pp. 1836–1840. 10.1007/0-306-48380-7_3459

Haselton, A.T., Yin, C.-M., Stoffolano, J.G., 2006. The effects of *Calliphora vomitoria* Tachykinin-I and the FMRFamide-related peptide Perisulfakinin on female *Phormia regina* crop contractions, in vitro. J Insect Physiol 52, 436–441. 10.1016/j.jinsphys.2005.12.003

Heath, A., Levot, G., 2015. Parasiticide resistance in flies, lice and ticks in New Zealand and Australia: mechanisms, prevalence and prevention. N Z Vet J 63, 199–210. 10.1080/00480169.2014.960500

Hermann, C., Yoshii, T., Dusik, V., Helfrich-Förster, C., 2012. Neuropeptide F immunoreactive clock neurons modify evening locomotor activity and free-running period in *Drosophila melanogaster*. Journal of Comparative Neurology 520, 970–987. 10.1002/cne.22742

Hobson RP, 1932. Studies on the nutrition of blow-fly larvae III. The liquefaction of muscle. 10.1242/jeb.8.2.109

Kannangara, J.R., Henstridge, M.A., Parsons, L.M., Kondo, S., Mirth, C.K., Warr, C.G., 2020. A New Role for Neuropeptide F Signaling in Controlling Developmental Timing and Body Size in *Drosophila melanogaster*. Genetics 216, 135–144. 10.1534/genetics.120.303475

Kapoor, S., Hickner, P. V., Dickey, A.N., Bailey, E., de Paula, L.C.B., Belikoff, E.J., Davis, R.J., Tandonnet, S., Canettieri, C.K., Bertone, M.A., Szpila, K., Hall, R.S., Young, N.D., Korhonen, P.K., Gasser, R.B., Perry, T., Jex, A.R., Bowles, V.M., Wiegmann, B.M., Torres, T.T., Anstead, C.A., Scott, M.J., 2025a. Comparative genomic analysis of necrophagous and parasitic subspecies of *Lucilia cuprina* (Diptera: Calliphoridae) provides important insight into their divergent biologies. Int J Parasitol. 10.1016/j.ijpara.2025.06.001

Kapoor, S., Yang, Y.T., Hall, R.N., Gasser, R.B., Bowles, V.M., Perry, T., Anstead, C.A., 2024. Complete Mitochondrial Genome for *Lucilia cuprina dorsalis* (Diptera: Calliphoridae) from the Northern Territory, Australia. Genes (Basel) 15, 506. 10.3390/genes15040506

Kapoor, S., Young, N.D., Jex, A.R., Baxter, S., Yang, Y.T., Davies, A.J., Armstrong, M.N., Whiteside, T.L., Batterham, P., Gasser, R.B., Bowles, V.M., Perry, T., Anstead, C.A., 2025b. Population structure, gene flow and genetic diversity of sheep blowfly (*Lucilia cuprina dorsalis*) in Australia. BMC Genomics 26. 10.1186/s12864-025-11852-y

Lee, S., Kim, Y.J., Jones, W.D., 2017. Central peptidergic modulation of peripheral olfactory responses. BMC Biol 15, 1. 10.1186/s12915-017-0374-6

Lennox, F.G., 1940. Distribution of Ammonia in Larvæ of *Lucilia cuprina*. Nature 146, 268–268. 10.1038/146268a0

Li, F., Wantuch, H.A., Linger, R.J., Belikoff, E.J., Scott, M.J., 2014. Transgenic sexing system for genetic control of the Australian sheep blow fly *Lucilia cuprina*. Insect Biochem Mol Biol 51, 80–88. 10.1016/j.ibmb.2014.06.001

Liu, B., Fu, D., Ning, H., Tang, M., Chen, H., 2021. Identification of the short neuropeptide f and short neuropeptide f receptor genes and their roles of food intake in dendroctonus armandi. Insects 12. 10.3390/insects12090844

Liu, W., Ganguly, A., Huang, J., Wang, Y., Ni, J.D., Gurav, A.S., Aguilar, M.A., Montell, C., 2019. Neuropeptide F regulates courtship in *Drosophila* through a male-specific neuronal circuit. Elife 8. 10.7554/eLife.49574

Liu, W., Longnecker, M., Tarone, A.M., Tomberlin, J.K., 2016. Responses of *Lucilia* sericata (Diptera: Calliphoridae) to compounds from microbial decomposition of larval resources. Anim Behav 115, 217–225. 10.1016/j.anbehav.2016.03.022

Love, M.I., Huber, W., Anders, S., 2014. Moderated estimation of fold change and dispersion for RNA-seq data with DESeq2. Genome Biol 15. 10.1186/s13059-014-0550-8

Martin, C., Minchilli, D., Francis, F., Verheggen, F., 2020. Behavioral and Electrophysiological Responses of the Fringed Larder Beetle *Dermestes frischii* to the Smell of a Cadaver at Different Decomposition Stages. Insects 11, 238. 10.3390/insects11040238

Matsushima, A., Yokotani, S., Lui, X., Sumida, K., Honda, T., Sato, S., Kaneki, A., Takeda, Y., Chuman, Y., Ozaki, M., Asai, D., Nose, T., Onoue, H., Ito, Y., Tominaga, Y., Shimohigashi, Y., Shimohigashi, M., 2003. Molecular cloning and circadian expression profile of insect neuropeptide PDF in black blowfly, *Phormia regina*. Int J Pept Res Ther 10, 419–430. 10.1007/s10989-004-2396-5

Mouga, D., Gaedke, A., 2017. Diptera survey in human corpses in the north of the state of Santa Catarina, Brazil. Acta Biológica Catarinense 4. 10.21726/abc.v4i1.359

Nasser, M.G., Hosni, E.M., Kenawy, M.A., Alharbi, S.A., Almoallim, H.S., Rady, M.H., Merdan, B.A., Pont, A.C., Al-Ashaal, S.A., 2021. Evolutionary profile of the family Calliphoridae, with notes on the origin of myiasis. Saudi J Biol Sci 28, 2056–2066. 10.1016/j.sjbs.2021.01.032

Nelson, L.A., Lambkin, C.L., Batterham, P., Wallman, J.F., Dowton, M., Whiting, M.F., Yeates, D.K., Cameron, S.L., 2012. Beyond barcoding: A mitochondrial genomics approach to molecular phylogenetics and diagnostics of blowflies (Diptera: Calliphoridae). Gene 511, 131–142. 10.1016/j.gene.2012.09.103

Post, Y., Paululat, A., 2018. Muscle Function Assessment Using a *Drosophila* Larvae Crawling Assay. Bio Protoc 8. 10.21769/BioProtoc.2933

Pu, Y., Zhang, Yiwen, Zhang, Yan, Shen, P., 2018. Two Drosophila Neuropeptide Y-like Neurons Define a Reward Module for Transforming Appetitive Odor Representations to Motivation. Sci Rep 8, 11658. 10.1038/s41598-018-30113-5

Sanford, M.R., 2017. Insects and associated arthropods analyzed during medicolegal death investigations in Harris County, Texas, USA: January 2013-April 2016. PLoS One 12, e0179404. 10.1371/journal.pone.0179404

Scherkenbeck, J., Zdobinsky, T., 2009. Insect neuropeptides: Structures, chemical modifications and potential for insect control. Bioorg Med Chem 17, 4071–4084. 10.1016/j.bmc.2008.12.061

Schoofs, L., De Loof, A., Van Hiel, M.B., 2017. Neuropeptides as Regulators of Behavior in Insects. Annu Rev Entomol 62, 35–52. 10.1146/annurev-ento-031616-035500

Setzu, M., Biolchini, M., Lilliu, A., Manca, M., Muroni, P., Poddighe, S., Bass, C., Angioy, A.M., Nichols, R., 2012. Neuropeptide F peptides act through unique signaling pathways to affect cardiac activity. Peptides (N.Y.) 33, 230–239. 10.1016/j.peptides.2012.01.005

Stone, B.L., Thummel, C.S., 1993. The Drosophila 78C early late puff contains E78, an ecdysone-inducible gene that encodes a novel member of the nuclear hormone receptor superfamily. Cell 75, 307–320. 10.1016/0092-8674(93)80072-M

Van Wielendaele, P., Wynant, N., Dillen, S., Zels, S., Badisco, L., Vanden Broeck, J., 2013. Neuropeptide F regulates male reproductive processes in the desert locust, *Schistocerca gregaria*. Insect Biochem Mol Biol 43, 252–259. 10.1016/j.ibmb.2012.12.004

Waterhouse, D., Paramonovo, S., 1950. The Status of the Two Species of Lucilia (Diptera, Calliphoridae) Attacking Sheep in Austhalia. Aust J Biol Sci 3, 310–336.

Williams, M.J., Akram, M., Barkauskaite, D., Patil, S., Kotsidou, E., Kheder, S., Vitale, G., Filaferro, M., Blemings, S.W., Maestri, G., Hazim, N., Vergoni, A.V., Schiöth, H.B., 2020. CCAP regulates feeding behavior via the NPF pathway in *Drosophila* adults. Proceedings of the National Academy of Sciences 117, 7401–7408. 10.1073/pnas.1914037117

Williamson, M.E., Yan, Y., Scott, M.J., 2021. Conditional knockdown of transformer in sheep blow fly suggests a role in repression of dosage compensation and potential for population suppression. PLoS Genet 17, e1009792. 10.1371/journal.pgen.1009792

Wu, Q., Wen, T., Lee, G., Park, J.H., Cai, H.N., Shen, P., 2003. Developmental Control of Foraging and Social Behavior by the Drosophila Neuropeptide Y-like System. Neuron 39, 147–161. 10.1016/S0896-6273(03)00396-9

Wu, T., Manogaran, A.L., Beauchamp, J.M., Waring, G.L., 2010. Drosophila vitelline membrane assembly: A critical role for an evolutionarily conserved cysteine in the “VM domain” of sV23. Dev Biol 347, 360–368. 10.1016/j.ydbio.2010.08.037

Wulff, J.P., Hickner, P. V., Watson, D.W., Denning, S.S., Belikoff, E.J., Scott, M.J., 2024. Antennal transcriptome analysis reveals sensory receptors potentially associated with host detection in the livestock pest *Lucilia cuprina*. Parasit Vectors 17, 308. 10.1186/s13071-024-06391-6

Wulff, J.P., Laminack, R.K., Scott, M.J., 2025. Genetic and behavioral analyses suggest that larval and adult stages of *Lucilia cuprina* employ different sensory systems to detect rotten beef. Parasit Vectors 18, 270. 10.1186/s13071-025-06804-0

Xie, H.-Y., Linghu, J.-H., Feng, J.-W., Yu, M.-Q., Xu, H.-M., Zhu, F., Yi, T.-C., Chen, X.-S., Smagghe, G., Liu, T.-X., Gui, S.-H., 2025. The NPF receptor regulates larval foraging behavior and female adult oviposition-site choice in *Spodoptera frugiperda*. Entomologia Generalis 45, 537–545. 10.1127/entomologia/2025/2991

Yoon, K.A., Kim, W.J., Cho, H., Yoon, H., Ahn, N.H., Lee, B.H., Lee, S.H., 2022. Characterization of anti-microbial peptides and proteins from maggots of Calliphoridae and Sarcophagidae fly species (Diptera). Comparative Biochemistry and Physiology Part - C: Toxicology and Pharmacology 259. 10.1016/j.cbpc.2022.109390

Zhu, J.J., Chaudhury, M.F., Tangtrakulwanich, K., Skoda, S.R., 2013. Identification of Oviposition Attractants of the Secondary Screwworm, *Cochliomyia macellaria (F.)* Released from Rotten Chicken Liver. J Chem Ecol 39, 1407–1414. 10.1007/s10886-013-0359-z

