## Supplementary material for "Neuropeptide F regulates adult female response to diet, larval locomotion and several larval physiological processes in *Lucilia cuprina cuprina*": MCI obtained from 11 replicate experiments was 0.49 (Table S1)

---

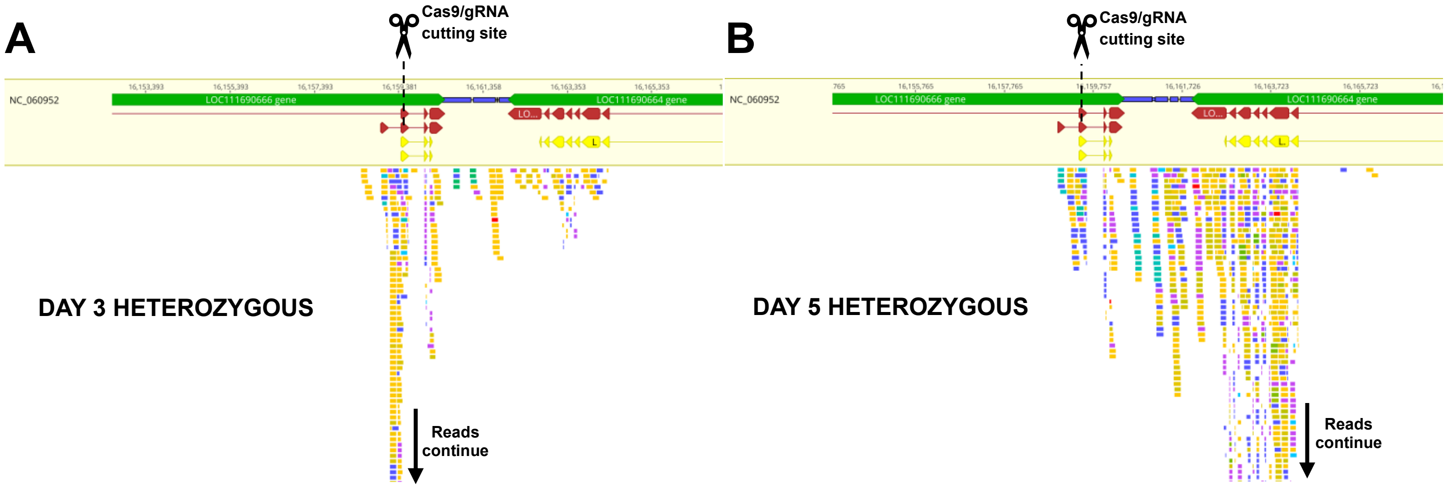

**Figure continued in the next page**

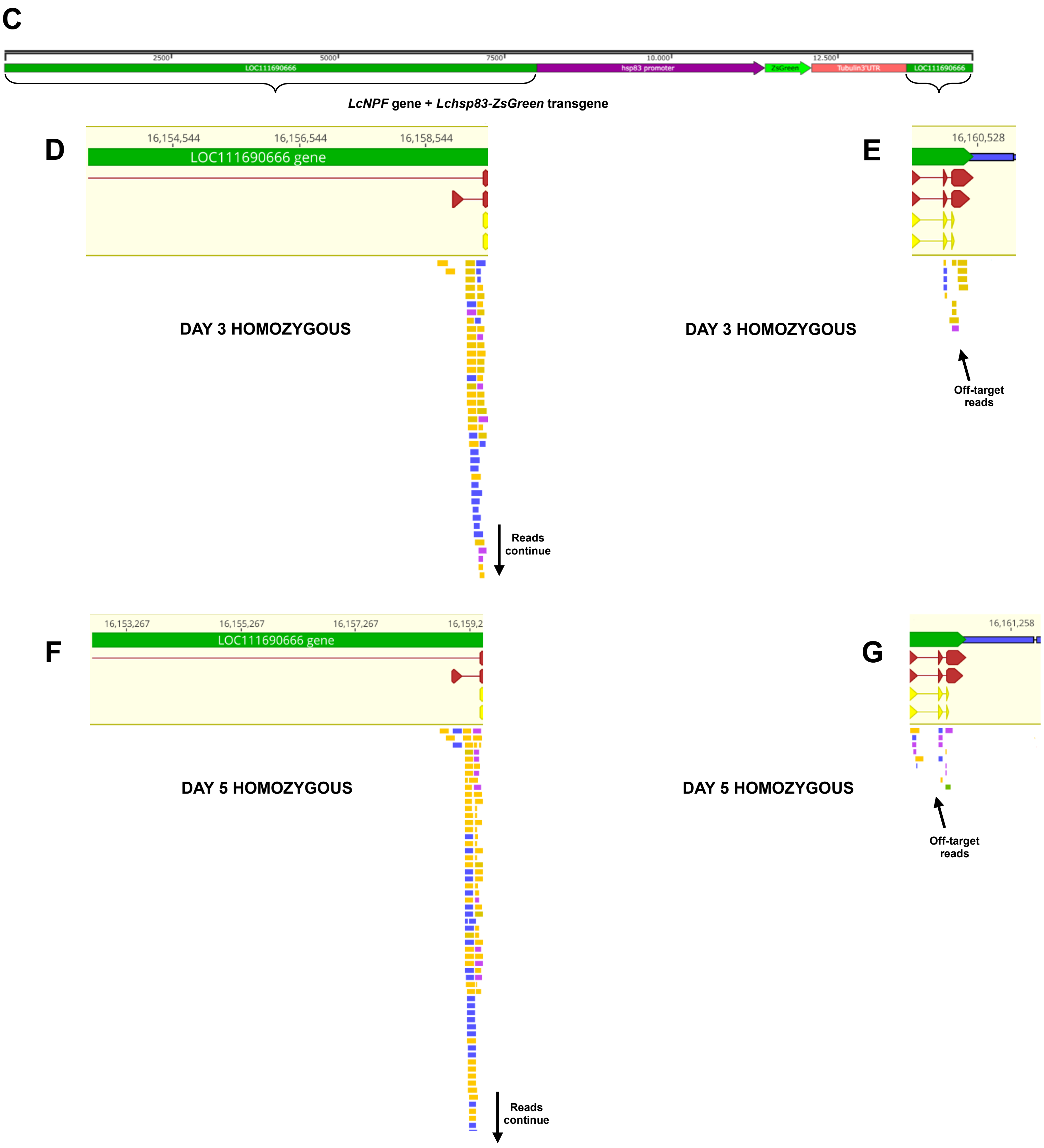

**Figure S1.** Mapping of RNA-Seq reads to the *LcNPF* gene region in heterozygous and homozygous *LcNPF* mutants. **A** and **B** show read alignments in heterozygous late L2 and early L3 larvae (days 3 and 5 respectively). In addition, the site of insertion of the *Lchsp83-ZsGreen* transgene is shown as the Cas9/gRNA cutting site in both quadrants. **C**. *LcNPF* gene + *Lchsp83-ZsGreen* transgene DNA map. **D** and **E** show read alignments in homozygous larvae on day 3, upstream and downstream the insert respectively. **F** and **G** present the alignments in homozygous larvae on day 5, upstream and downstream the insert respectively. Note that only the forward and reverse sequence of an off-target read was mapping the *LcNPF* gene in homozygous larvae following the *Lchsp83-ZsGreen* transgene: ACAAGCCGAT+ACGTGTGTCA and GACACCACTT+TCAGGCTTGT (reverse orientation).

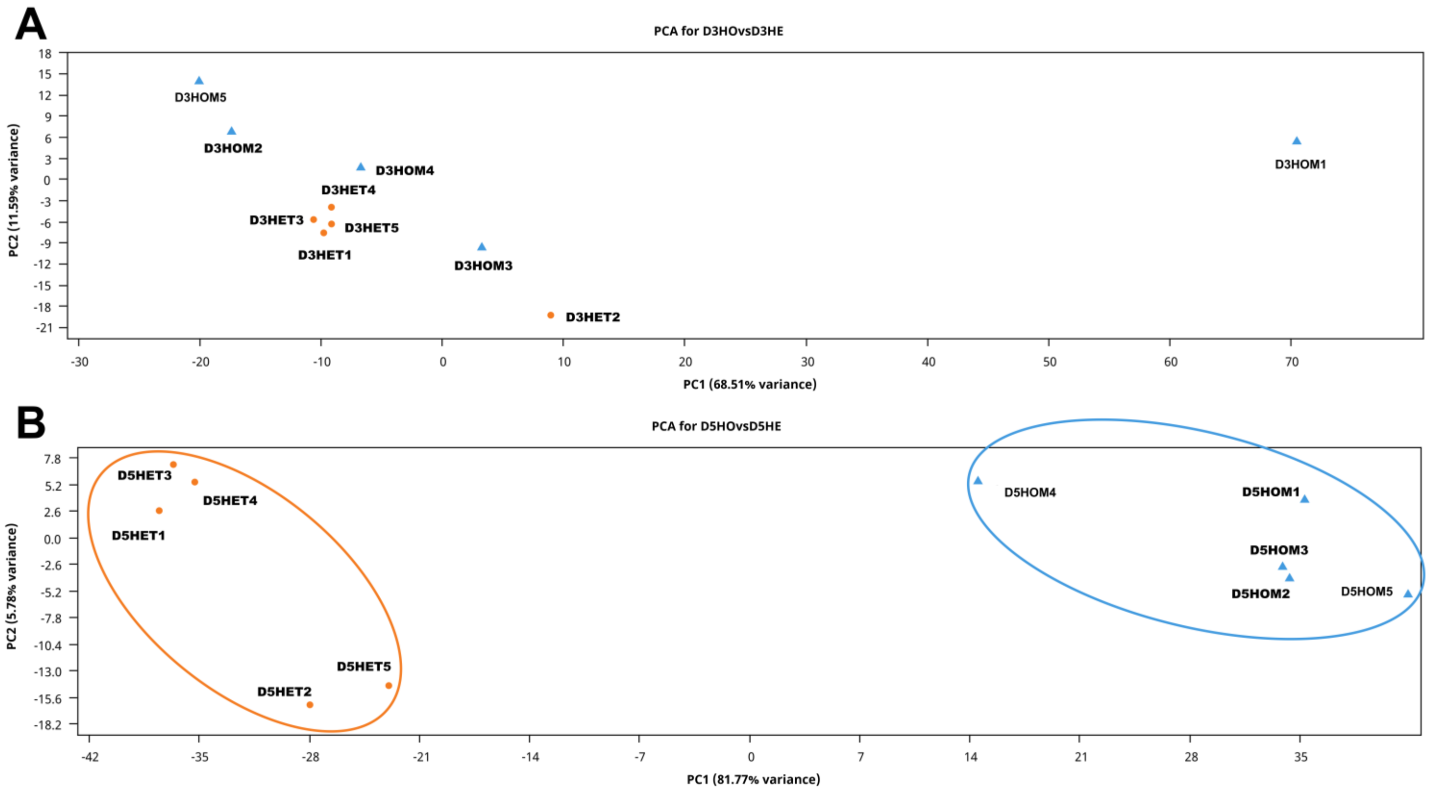

**Figure S2.** Principal Component Analysis (PCA) of RNA-Seq data. **A**. Libraries completed using heterozygous (HET) and homozygous (HOM) larvae collected at day 3 (D3) after egg hatching. **B**. Libraries completed using HET and HOM larvae collected on day 5 (D5) after egg hatching. Note that libraries of different genotypes, *i.e*. HET and HOM, were only separated on day 5, notably, with PC1 including 81.77% of the total variance of the data (highlighted in orange and light blue, respectively). Before the PCA analysis, read counts were normalized using DESeq2 median of ratios normalization method.

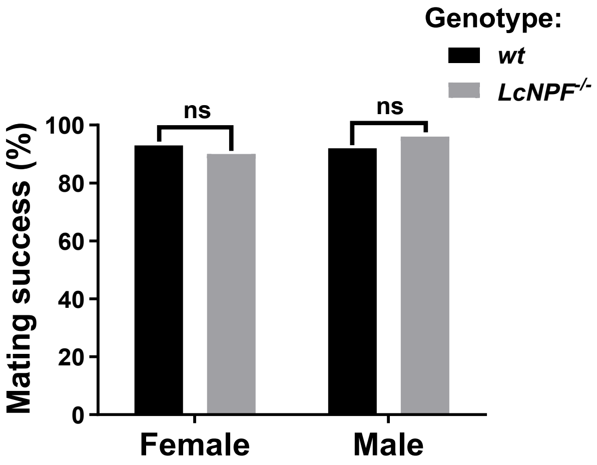

**Figure S3**. *LcNPF*$^{-/-}$ strain fertility and mating fitness evaluation. Fertility of both sexes was normal when compared to *wt* flies. Abbreviations: *LcNPF*$^{-/-}$ = NPF null mutated larvae; *wt* = wild-type.

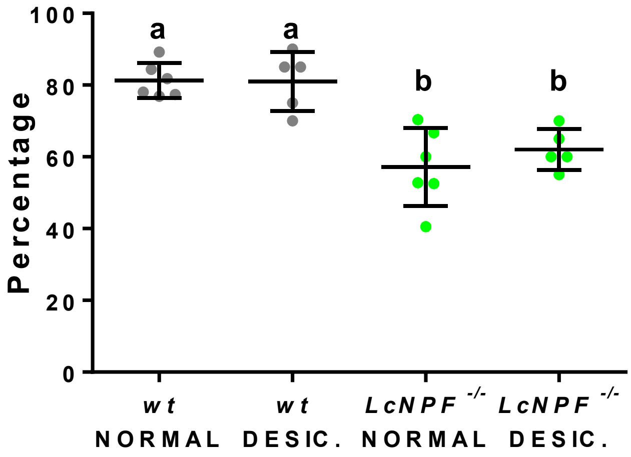

**Figure S4**. Proportion of wild-type (*wt*) and *LcNPF*$^{-/-}$ mutant embryos that complete development under normal and desiccation conditions. Under normal conditions, the average egg hatching rate was 81% for the LA07 *wt* and 57% for the *LcNPF*$^{-/-}$ strain. Following a desiccation treatment, hatching rates remained similar, with 81% for LA07 *wt* and 62% for *LcNPF*$^{-/-}$. Statistical analysis revealed a significant difference (different lowercase letters) in hatching rates (in percentages) between genotypes, whereas no significant effect of desiccation treatment was observed. Statistical analysis was competed using ANOVA followed by the Tukey's HSD test for multiple comparisons (n = 5-6 replicates, with 20–30 eggs per replicate). Abbreviations: desic. = desiccation; *LcNPF*$^{-/-}$ = NPF null mutated larvae; *wt* = wild-type.

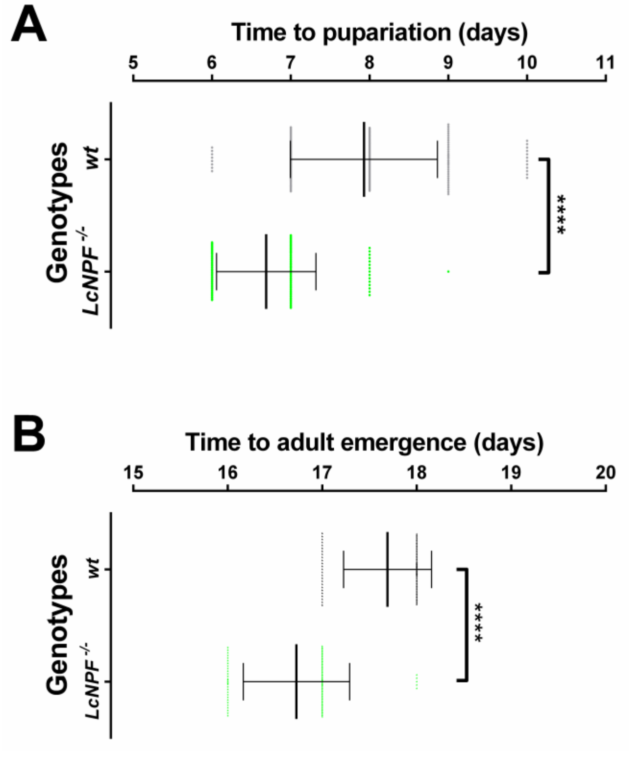

**Figure S5**. **A**. Time to pupariation for *wt* and *LcNPF*$^{-/-}$ mutant individuals. The time to pupariation after egg hatching was statistically significant between individuals obtained from the LA07 *wt* and *LcNPF*$^{-/-}$ colony. **B**. Time to adult emergence for *wt* and *LcNPF* mutants. The time to adult emergence was statistically significant between individuals obtained from the LA07 *wt* and *LcNPF*$^{-/-}$ colonies. A total of 100 wandering L3 larvae (**A**) or pupae (**B**) were analyzed for each group. Dots represent single larvae or adults, and the mean time to puparation (**A**) or adult emergence (**B**) for each group ± (standard deviation) SD was plotted. A Mann Whitney test was used for statistical analysis of both experiments. Four asterisks correspond to $P < 0.0001$. Abbreviations: *LcNPF*$^{-/-}$ = NPF null mutated larvae; *wt* = wild-type.

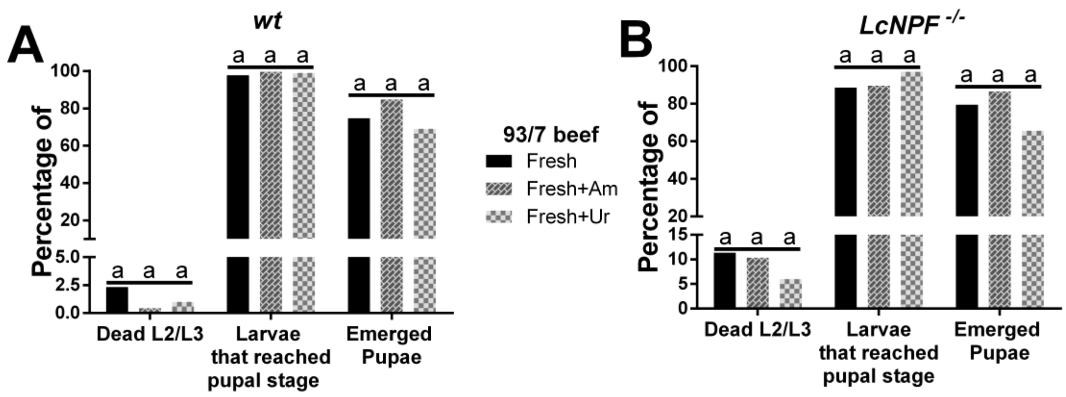

**Figure S6**. Survival of *wt* and *LcNPF*$^{-/-}$ larvae reared on fresh beef or beef supplemented with either ammonium or urea. Wild-type (**A**) or *LcNPF*$^{-/-}$ larvae (**B**) were reared on 93/7% meat alone or mixed with ammonium hydroxide or urea (both 1% final concentration). The percentage (%) of dead larvae during the L2/L3 stages and larvae that reached the pupal stage were calculated from the total (T, see Table S3) resulting from the sum of both. The % of emerged pupae were calculated from the number of adults obtained over the number of pupae. Abbreviations: AVG = average; Am = ammonium hydroxide; fresh or F = fresh beef; rotten or R = rotten beef; T = total; Ur = urea; *LcNPF*$^{-/-}$ = NPF null mutated larvae; *wt* = wild-type; % Pup. Em = percentage of pupae emergence. Median's groups were compared using Kruskal-Wallis test followed by Dunn's test correction for multiple comparisons, and same lowercase letters means non statistically significant.

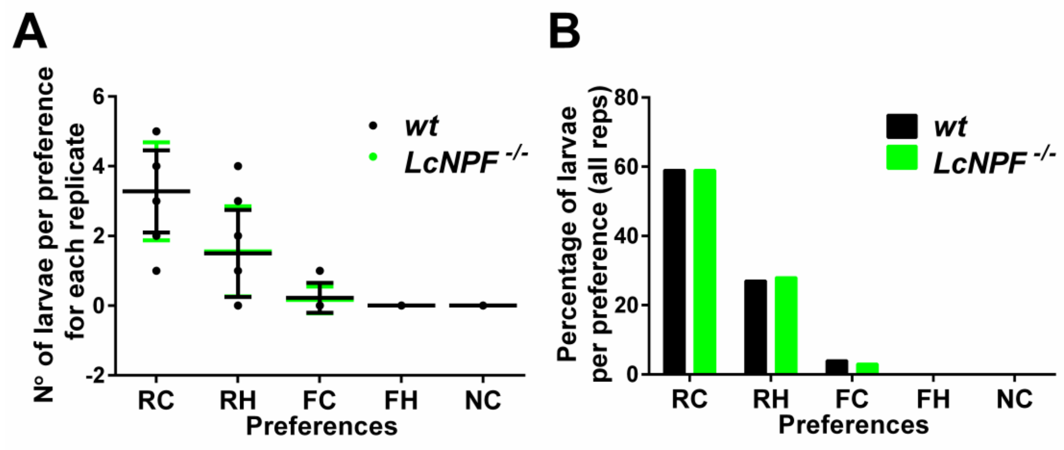

**Figure S7**. Larval diet and temperature preference test, using a mix of late-L2 + early-L3 larvae. Larvae were given a choice of fresh and rotting 93/7% beef at room temperature or 37°C as described previously (Wulff *et al*. 2025). **A**. The number of larvae choosing each condition with the average number of larvae is shown. **B**. The percentage of larvae that have chosen each condition for each genotype calculated on the total number of larvae tested. The larval preference was not statistically significant when comparing larvae obtained from the LA07 *wt* and *LcNPF*$^{-/-}$ colonies. Abbreviations: FC = fresh cold beef; FH = fresh hot beef; RC = rotten cold beef; RH = rotten hot beef; NC = non-choice; *LcNPF*$^{-/-}$ = NPF null mutated larvae; *wt* = wild-type.

### Supplementary Tables

**Table S1.** Homozygous *LcNPF*$^{-/-}$ males are competitive with wild type males for mating with wild type females.

| Bottle | Number females alive after 16h with males | Number fertile females | Number females mated with wild type males | Number females mated with *LcNPF*$^{-/-}$ males | MCI |
|---|---|---|---|---|---|
| A | 9 | 6 | 5 | 1 | 0.17 |
| B | 9 | 7 | 5 | 2 | 0.29 |
| C | 10 | 9 | 4 | 5 | 0.55 |
| D | 10 | 10 | 4 | 6 | 0.6 |
| E | 9 | 8 | 6 | 2 | 0.25 |
| F | 9 | 8 | 3 | 5 | 0.62 |
| G | 8 | 7 | 2 | 5 | 0.71 |
| H | 8 | 8 | 3 | 5 | 0.62 |
| I | 6 | 4 | 1 | 3 | 0.75 |
| J | 9 | 8 | 4 | 4 | 0.5 |
| K | 9 | 8 | 5 | 3 | 0.38 |
| Overall | 96 | 83 | 42 | 41 | 0.494 |

Abbreviations: MCI = mating competitiveness index.

**Table S2**. NC-LA07 *wt* and *LcNPF*$^{-/-}$ larvae survival, reared in fresh and rotten lean-fat 73/27%.

| | Dead L2/L3 | % | Pupae | % | T | Adult | % Pup. Em |
|---|---|---|---|---|---|---|---|
| *wt* - Fresh_1 | 18 | 8.7 | 188 | 91.3 | 206 | 66 | 35.1 |
| *wt* - Fresh_2 | 21 | 10.5 | 179 | 89.5 | 200 | 63 | 35.2 |
| *wt* - Fresh_3 | 30 | 13.4 | 194 | 86.6 | 224 | 96 | 49.5 |
| **AVG** | | **11** | | **89** | | | **40** |
| *wt* - Rotten_1 | 162 | 65.9 | 84 | 34.1 | 246 | 21 | 25.0 |
| *wt* - Rotten_2 | 141 | 87.0 | 21 | 13.0 | 162 | 0 | 0.0 |
| *wt* - Rotten_3 | 168 | 93.3 | 12 | 6.7 | 180 | 3 | 25.0 |
| **AVG** | | **82** | | **18** | | | **17** |
| *LcNPF*$^{-/-}$ - Fresh_1 | 54 | 25.6 | 157 | 74.4 | 211 | 45 | 28.7 |
| *LcNPF*$^{-/-}$ - Fresh_2 | 129 | 57.3 | 96 | 42.7 | 225 | 6 | 6.3 |
| *LcNPF*$^{-/-}$ - Fresh_3 | 93 | 43.1 | 123 | 56.9 | 216 | 21 | 17.1 |
| **AVG** | | **42** | | **58** | | | **17** |
| *LcNPF*$^{-/-}$ - Rotten_1 | 192 | 100.0 | 0 | 0.0 | 192 | 0 | - |
| *LcNPF*$^{-/-}$ - Rotten_2 | 216 | 100.0 | 0 | 0.0 | 216 | 0 | - |
| *LcNPF*$^{-/-}$ - Rotten_3 | 240 | 100.0 | 0 | 0.0 | 240 | 0 | - |
| **AVG** | | **100** | | **0** | | | |

The percentage (%) of dead larvae during the L2/L3 stages and larvae that reached the pupal stage were calculated from the total (T) resulting from the sum of both. The % of emerged pupae were calculated from the number of adults obtained over the number of pupae. Abbreviations: AVG = average; *LcNPF*$^{-/-}$ = NPF-null mutated larvae; *wt* = wild-type; % Pup. Em = percentage of pupae emergence.

**Table S3**. NC-LA07 *wt* and *LcNPF*$^{-/-}$ larvae survival, reared in lean-fat 93/7% ground beef under different conditions.

| | Dead L2/L3 | % | Pupae | % | T | Adult | % Pup. Em |
|---|---|---|---|---|---|---|---|
| *wt* - Fresh_1 | 8 | 3 | 274 | 97 | 282 | 159 | 58 |
| *wt* - Fresh_2 | 5 | 2 | 199 | 98 | 204 | 158 | 79 |
| *wt* - Fresh_3 | 7 | 2 | 312 | 98 | 319 | 270 | 87 |
| **AVG** | | **2** | | **98** | | | **75** |
| *wt* - Fresh+Am_1 | 1 | 0.3 | 353 | 100 | 354 | 337 | 95 |
| *wt* - Fresh+Am_2 | 0 | 0.0 | 370 | 100 | 370 | 315 | 85 |
| *wt* - Fresh+Am_3 | 4 | 1.0 | 388 | 99 | 392 | 287 | 74 |
| **AVG** | | **0** | | **100** | | | **85** |
| *wt* - Fresh+Ur_1 | 2 | 1 | 285 | 99 | 287 | 196 | 69 |
| *wt* - Fresh+Ur_2 | 4 | 1 | 302 | 99 | 306 | 202 | 67 |
| *wt* - Fresh+Ur_3 | 3 | 1 | 278 | 99 | 281 | 198 | 71 |
| **AVG** | | **1** | | **99** | | | **69** |
| *wt* - Rotten_1 | 196 | 47 | 217 | 53 | 413 | 120 | 55 |
| *wt* - Rotten_2 | 200 | 46 | 234 | 54 | 434 | 167 | 72 |
| *wt* - Rotten_3 | 166 | 46 | 196 | 54 | 362 | 144 | 73 |
| **AVG** | | **46** | | **54** | | | **67** |
| *LcNPF*$^{-/-}$ - Fresh_1 | 8 | 4 | 200 | 96 | 208 | 117 | 58 |
| *LcNPF*$^{-/-}$ - Fresh_2 | 40 | 17 | 204 | 83 | 244 | 167 | 82 |
| *LcNPF*$^{-/-}$ - Fresh_3 | 30 | 11 | 249 | 89 | 279 | 144 | 58 |
| *LcNPF*$^{-/-}$ - Fresh_4 | 34 | 11 | 271 | 89 | 305 | 252 | 93 |
| *LcNPF*$^{-/-}$ - Fresh_5 | 35 | 13 | 242 | 87 | 277 | 230 | 95 |
| *LcNPF*$^{-/-}$ - Fresh_6 | 48 | 12 | 363 | 88 | 411 | 331 | 91 |
| **AVG** | | **10** | | **90** | | | **66** |
| *LcNPF*$^{-/-}$ - Fresh+Am_1 | 56 | 12 | 396 | 88 | 452 | 367 | 93 |
| *LcNPF*$^{-/-}$ - Fresh+Am_2 | 35 | 9 | 367 | 91 | 402 | 330 | 90 |
| *LcNPF*$^{-/-}$ - Fresh+Am_3 | 38 | 10 | 342 | 90 | 380 | 264 | 77 |
| **AVG** | | **10** | | **90** | | | **87** |
| *LcNPF*$^{-/-}$ - Fresh+Ur_1 | 19 | 6 | 320 | 94 | 339 | 206 | 64 |
| *LcNPF-*$^{/-}$ - Fresh+Ur_2 | 21 | 7 | 289 | 93 | 310 | 196 | 68 |
| *LcNPF*$^{-/-}$ - Fresh+Ur_3 | 16 | 5 | 291 | 95 | 307 | 190 | 65 |
| **AVG** | | **6** | | **94** | | | **66** |
| *LcNPF*$^{-/-}$ - Rotten_1 | 324 | 89 | 40 | 11 | 364 | 5 | 13 |
| *LcNPF*$^{-/-}$ - Rotten_2 | 267 | 90 | 30 | 10 | 297 | 3 | 10 |
| *LcNPF*$^{-/-}$ - Rotten_3 | 300 | 90 | 34 | 10 | 334 | 6 | 18 |
| **AVG** | | **90** | | **10** | | | **13** |

The percentage (%) of dead larvae during the L2/L3 stages and larvae that reached the pupal stage were calculated from the total (T) resulting from the sum of both. The % of emerged pupae were calculated from the number of adults obtained over the number of pupae. Abbreviations: AVG = average; Am = ammonium hydroxide; T = total; Ur = urea; *LcNPF*$^{-/-}$ = NPF-null mutated larvae; *wt* = wild-type; % Pup. Em = percentage of pupae emergence.

**Table S4**. NC-LA07 *wt* and *LcNPF*$^{-/-}$ adult female olfaction assay.

| Time | ***wt* FRESH** | | | | | | | | | | | | | | | | | | | | | | | | |
|---|---|---|---|---|---|---|---|---|---|---|---|---|---|---|---|---|---|---|---|---|---|---|---|---|---|
| 15min | 0 | 0 | 1 | 0 | 0 | 0 | 0 | 0 | 0 | 0 | 0 | 0 | 0 | 0 | 0 | 0 | 0 | 0 | 0 | 0 | 0 | 0 | 0 | 0 | 0 |
| 30min | 0 | 0 | 1 | 1 | 0 | 0 | 0 | 0 | 0 | 0 | 0 | 0 | 0 | 0 | 0 | 0 | 0 | 0 | 0 | 0 | 0 | 1 | 0 | 0 | 0 |
| 45min | 0 | 0 | 1 | 1 | 0 | 0 | 0 | 0 | 0 | 0 | 0 | 0 | 0 | 0 | 0 | 0 | 0 | 0 | 0 | 0 | 0 | 1 | 0 | 1 | 0 |
| 1 h | 0 | 2 | 1 | 2 | 0 | 0 | 0 | 0 | 0 | 1 | 0 | 0 | 0 | 0 | 1 | 0 | 0 | 0 | 0 | 0 | 0 | 1 | 0 | 1 | 0 |
| 2 h | 0 | 2 | 1 | 5 | 4 | 0 | 0 | 1 | 0 | 2 | 0 | 0 | 1 | 0 | 2 | 0 | 0 | 0 | 0 | 0 | 0 | 1 | 2 | 1 | 1 |
| 3 h | 0 | 2 | 1 | 5 | 4 | 0 | 0 | 1 | 1 | 3 | 0 | 0 | 1 | 1 | 3 | 0 | 0 | 0 | 0 | 0 | 0 | 1 | 2 | 1 | 1 |
| 4 h | 0 | 3 | 1 | 6 | 4 | 0 | 0 | 2 | 1 | 3 | 0 | 0 | 2 | 1 | 3 | 0 | 0 | 0 | 0 | 0 | 0 | 1 | 2 | 1 | 1 |
| O/N | 0 | 3 | 1 | 6 | 4 | 3 | 0 | 3 | 1 | 3 | 3 | 0 | 3 | 1 | 3 | 0 | 1 | 0 | 1 | 1 | 0 | 2 | 3 | 1 | 1 |

| Time | ***wt* ROTTEN** | | | | | | | | | | | | | | | | | | | | | | | | |
|---|---|---|---|---|---|---|---|---|---|---|---|---|---|---|---|---|---|---|---|---|---|---|---|---|---|
| 15min | 0 | 1 | 0 | 0 | 0 | 0 | 0 | 0 | 0 | 0 | 0 | 0 | 0 | 0 | 0 | 0 | 0 | 0 | 0 | 0 | 0 | 0 | 0 | 1 | 0 |
| 30min | 0 | 1 | 0 | 0 | 0 | 0 | 0 | 0 | 0 | 0 | 0 | 0 | 0 | 0 | 0 | 0 | 0 | 2 | 4 | 3 | 0 | 1 | 0 | 1 | 0 |
| 45min | 0 | 1 | 0 | 1 | 0 | 0 | 0 | 2 | 2 | 1 | 0 | 0 | 2 | 2 | 1 | 2 | 0 | 4 | 5 | 5 | 0 | 1 | 0 | 3 | 0 |
| 1 h | 0 | 3 | 0 | 1 | 0 | 0 | 0 | 2 | 3 | 1 | 0 | 0 | 2 | 3 | 1 | 3 | 0 | 4 | 5 | 5 | 0 | 1 | 0 | 3 | 3 |
| 2 h | 2 | 6 | 2 | 2 | 3 | 0 | 2 | 2 | 5 | 4 | 0 | 2 | 2 | 5 | 4 | 4 | 1 | 6 | 6 | 5 | 6 | 3 | 2 | 6 | 5 |
| 3 h | 3 | 6 | 2 | 3 | 5 | 2 | 6 | 2 | 5 | 4 | 2 | 6 | 2 | 5 | 4 | 5 | 1 | 6 | 6 | 6 | 7 | 4 | 3 | 8 | 6 |
| 4 h | 3 | 6 | 3 | 4 | 5 | 2 | 6 | 2 | 6 | 4 | 2 | 6 | 2 | 6 | 4 | 6 | 3 | 9 | 6 | 6 | 7 | 4 | 4 | 8 | 6 |
| O/N | 3 | 6 | 4 | 4 | 5 | 2 | 6 | 2 | 6 | 4 | 2 | 6 | 2 | 6 | 4 | 8 | 7 | 9 | 6 | 7 | 8 | 5 | 5 | 8 | 7 |

| Time | ***LcNPF*$^{-/-}$ FRESH** | | | | | | | | | | | | | | | | | | | | | | | | |
|---|---|---|---|---|---|---|---|---|---|---|---|---|---|---|---|---|---|---|---|---|---|---|---|---|---|
| 15min | 0 | 0 | 0 | 0 | 0 | 0 | 0 | 0 | 0 | 0 | 0 | 0 | 0 | 0 | 0 | 0 | 2 | 0 | 0 | 0 | 0 | 0 | 0 | 0 | 0 |
| 30min | 0 | 0 | 1 | 0 | 0 | 0 | 0 | 0 | 0 | 0 | 0 | 0 | 0 | 0 | 0 | 0 | 2 | 2 | 1 | 0 | 0 | 0 | 0 | 0 | 0 |
| 45min | 0 | 0 | 1 | 0 | 1 | 0 | 0 | 0 | 0 | 0 | 0 | 0 | 0 | 0 | 0 | 0 | 2 | 2 | 1 | 1 | 0 | 0 | 0 | 0 | 0 |
| 1 h | 0 | 0 | 1 | 1 | 1 | 0 | 0 | 0 | 1 | 0 | 0 | 0 | 0 | 1 | 0 | 0 | 2 | 2 | 1 | 1 | 0 | 0 | 0 | 0 | 0 |
| 2 h | 1 | 0 | 3 | 3 | 2 | 2 | 0 | 1 | 1 | 0 | 2 | 0 | 1 | 1 | 0 | 1 | 3 | 3 | 4 | 1 | 2 | 0 | 3 | 2 | 3 |
| 3 h | 2 | 1 | 3 | 3 | 4 | 2 | 0 | 1 | 1 | 0 | 2 | 0 | 1 | 1 | 0 | 1 | 3 | 5 | 4 | 1 | 3 | 0 | 4 | 3 | 4 |
| 4 h | 3 | 2 | 4 | 3 | 4 | 3 | 0 | 1 | 1 | 0 | 3 | 0 | 1 | 1 | 0 | 1 | 3 | 5 | 4 | 1 | 4 | 0 | 5 | 3 | 4 |
| O/N | 3 | 3 | 4 | 3 | 5 | 4 | 1 | 2 | 4 | 0 | 4 | 1 | 2 | 4 | 0 | 4 | 4 | 6 | 4 | 2 | 4 | 1 | 6 | 4 | 5 |

| Time | ***LcNPF*$^{-/-}$ ROTTEN** | | | | | | | | | | | | | | | | | | | | | | | | |
|---|---|---|---|---|---|---|---|---|---|---|---|---|---|---|---|---|---|---|---|---|---|---|---|---|---|
| 15min | 0 | 0 | 0 | 0 | 0 | 0 | 0 | 0 | 0 | 0 | 0 | 0 | 0 | 0 | 0 | 0 | 0 | 0 | 0 | 0 | 0 | 0 | 0 | 0 | 0 |
| 30min | 0 | 0 | 0 | 0 | 0 | 0 | 0 | 0 | 0 | 0 | 0 | 0 | 0 | 0 | 0 | 0 | 0 | 0 | 0 | 0 | 0 | 0 | 0 | 0 | 0 |
| 45min | 0 | 0 | 0 | 0 | 0 | 0 | 0 | 0 | 0 | 0 | 0 | 0 | 0 | 0 | 0 | 0 | 0 | 0 | 0 | 0 | 0 | 0 | 1 | 0 | 0 |
| 1 h | 0 | 0 | 0 | 0 | 0 | 0 | 0 | 0 | 0 | 1 | 0 | 0 | 0 | 0 | 1 | 0 | 0 | 0 | 0 | 0 | 0 | 0 | 1 | 0 | 0 |
| 2 h | 1 | 2 | 0 | 0 | 0 | 0 | 1 | 2 | 1 | 3 | 0 | 1 | 2 | 1 | 3 | 1 | 0 | 0 | 1 | 0 | 0 | 2 | 2 | 1 | 1 |
| 3 h | 2 | 2 | 0 | 0 | 2 | 0 | 1 | 2 | 1 | 3 | 0 | 1 | 2 | 1 | 3 | 1 | 1 | 0 | 1 | 2 | 0 | 5 | 3 | 4 | 4 |
| 4 h | 2 | 2 | 1 | 0 | 2 | 1 | 2 | 3 | 2 | 4 | 1 | 2 | 3 | 2 | 4 | 3 | 2 | 0 | 2 | 2 | 0 | 7 | 3 | 5 | 4 |
| O/N | 2 | 3 | 3 | 0 | 3 | 1 | 2 | 5 | 3 | 4 | 1 | 2 | 5 | 3 | 4 | 3 | 4 | 1 | 6 | 8 | 1 | 8 | 4 | 6 | 5 |

Numbers represent the number of adult females attracted to each type of meat (*i.e*. fresh or rotten lean-fat 93/7% ground beef) at different times. Abbreviations: h = hour, O/N = overnight.

**Table S5**. NC-LA07 *wt* adult female oviposition substrate preference.

| **Substrate** | **73/27 F** | **73/27 R** | **93/7 F** | **93/7 R** | **NC** | **TOTAL** |
|---|---|---|---|---|---|---|
| **Number of females** | 1 | 9 | 1 | 1 | 8 | 20 |
| **Percentage** | 5 | 45 | 5 | 5 | 40 | 100 |
| **Substrate** | **73/27 F** | **73/27 R** | **NC** | **TOTAL** | | |
| **Number of females** | 1 | 19 | 5 | 25 | | |
| **Percentage** | 4 | 76 | 20 | 100 | | |

Abbreviations: NC = non-choice; F = fresh; R = rotten.

**Table S6**. NC-LA07 *wt* and *LcNPF*$^{-/-}$ adult female oviposition substrate preference.

| | | | | | | | | | | | | | | | | | | | | | | | | | |
|---|---|---|---|---|---|---|---|---|---|---|---|---|---|---|---|---|---|---|---|---|---|---|---|---|---|
| ***wt*** | 0 | 1 | 1 | 1 | 1 | 1 | 1 | 1 | 1 | 1 | 1 | 1 | 1 | 1 | 1 | 1 | 1 | 1 | 1 | 1 | 2 | 2 | 2 | 2 | 2 |
| ***LcNPF*$^{-/-}$** | 0 | 0 | 0 | 0 | 0 | 0 | 0 | 0 | 0 | 0 | 0 | 0 | 0 | 0 | 0 | 0 | 0 | 0 | 0 | 1 | 1 | 1 | 2 | 2 | 2 |

Code number: 0, 1 and 2 represent females that showed a preference for fresh or rotten meat, or did not choose any option, respectively.
