## Supplementary material for "Neuropeptide F regulates adult female response to diet, larval locomotion and several larval physiological processes in *Lucilia cuprina cuprina*": in the supplementary method (SM) section SM1A-C

**A**

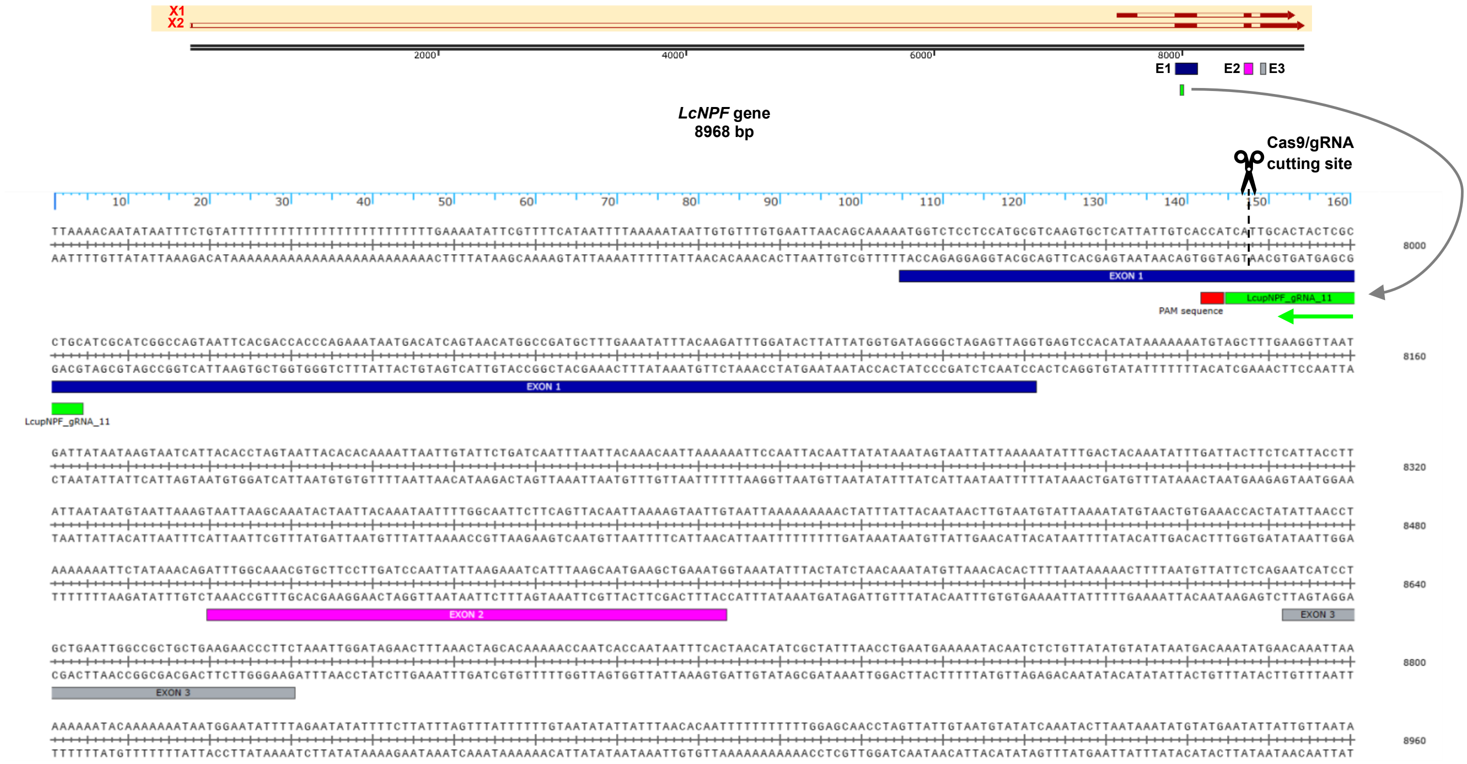


**B**

```
ATGGTCTCCTCCATGCGTCAAGTGCTCATTATTGTCACCATCATTGCACTACTCGCCTGCATCGCATCGGCCAGTAATTCACGACCACCCAGAAATAATGACATCAGTAACA
TGGCCGATGCTTTGAAATATTTACAAGATTTGGATACTTATTATGGTGATAGGGCTAGAGTTAGATTTGGCAAACGTGCTTCCTTGATCCAATTATTAAGAAATCATTTAAG
CAATGAAGCTGAAATGAATCATCCTGCTGAATTGGCCGCTGCTGAAGAACCCTTCTAA
```

**C**

**Cas9/gRNA cutting site**

MVSSMRQVLIIVTIIALLACIASASNSRPPRNNDISNMADALKYLQDLDTYYGDRA**RVRF**GKRASLIQLLRNHLSN
EAEMNHPAELAAAEEPF

Signal-Peptide
**Cleavage site**
Cleaved pre-active peptide
**Conserved motif**

**Supplementary method 1**. *Lucilia cuprina cuprina* Neuropeptide F gene [*LcNPF*), LOC111690666] gene and protein sequences: **A**. *LcNPF* gene, including gene variants X1 and X2. First, second and third exons were detailed in blue, purple and gray, respectively. The guide (g) RNA LcupNPF-gRNA-11 and PAM locations were detailed in green and red, respectively. The CRISPR/Cas9 cleavage site was highlighted with scissors and a dashed line. In addition, **B** and **C**

provide the *LcNPF* CDS and the pre-propeptide sequences, respectively. Peptide features are detailed by a color code below the pre-propeptide. The CRISPR/Cas9 cleavage site was highlighted with scissors and a dashed line.

**A**

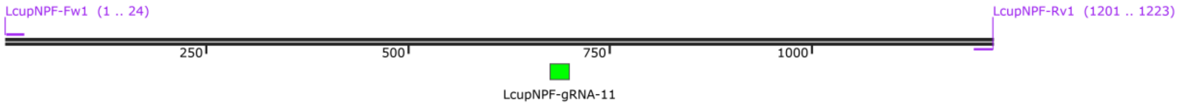


***LcNPF* gene fragment (for in vitro test)**
**1223 bp**

**B**

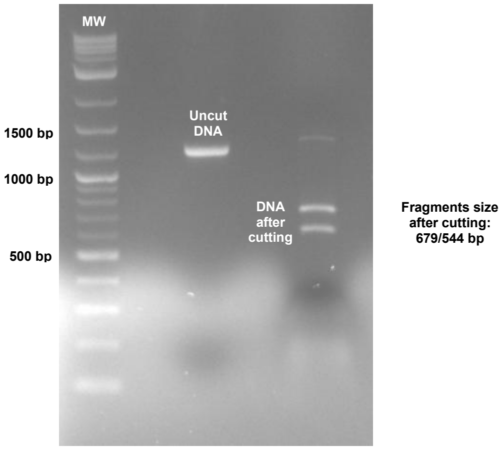


**Supplementary method 2**. Cutting efficiency of LcupNPF-gRNA-11 using a Cas9 in vitro assay: **A**. Amplification of the *LcNPF* DNA fragment to be used as a template in the CRISPR/Cas9 in vitro assay. The *LcNPF* fragment of 1223 base pairs (bp), flanking the LcupNPF-gRNA-11 location, was amplified by polymerase chain reaction (PCR) for 40 cycles, using as template *L. cuprina wt* genomic (g) DNA obtained from adult flies of mixed sexes, the Q5® High-Fidelity DNA Polymerase [Q5-Taq polymerase, New England Biolabs (NEB), Ipswich, MA, USA, Cat. #M0491S] and primers LcupNPF-Fw1: ACTGTAAGTGAGAGTGTGTGTGTT and LcupNPF-Rv1: ATTGGATCAAGGAAGCACGTTTG. The product was purified using the DNA Clean Kit (Zymo Research, Irvine, CA, USA, Cat. #D4004) following manufacturer's specifications and diluted to 20 ng/µl concentration. **B**. *LcNPF* CRISPR/Cas9 in vitro assays. The LcupNPF-gRNA-11 Cas9-crRNA [Integrated DNA Technologies (IDT), Coralville, IA, USA] was mixed with the Cas9-tracrRNA (IDT, Cat. #1072532) to generate the duplex crRNA::tracrRNA using an equal concentration of both RNAs. The mix was heated at 95 ºC for 5 min, cooled to ~15 ºC for 5 min and kept at -20 ºC until use. Subsequently, the Cas9 and gRNA duplex were mixed for a final concentration of 66.6 and 160 nM, respectively, and incubated for 20 min at 25°C to form the Cas9 ribonucleoprotein (RNP) complex. Following, the purified PCR product (*LcNPF* DNA fragment) obtained in **A** was added to the mix for a final concentration of 3 nM and incubated for 60 min at 25°C. The reaction (RXN) was ended by adding 1 µl of Proteinase K 10 µg/µl (Zymo Research, Cat. #D3001-2-20) and incubated for 10 min. The RXN mix was loaded into a 1.5% agarose gel in parallel to uncut gDNA (same DNA used as a template at the same concentration) and a mass weight (MW) ladder, electrophoresed at 60 volts for 90 min and visualize using a Gel Doc™ EZ System (Bio-Rad, Hercules, CA, USA). Two bands at approximately the expected size, *i.e*. 679 bp and 544 bp, confirmed the successful in vitro cleavage activity of LcupNPF-gRNA-11.

**A**

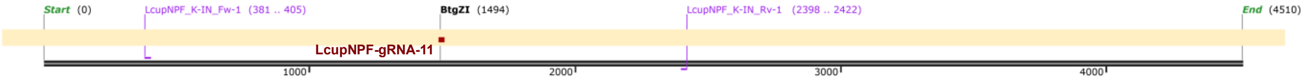


***LcNPF* gene fragment**
**4510 bp**

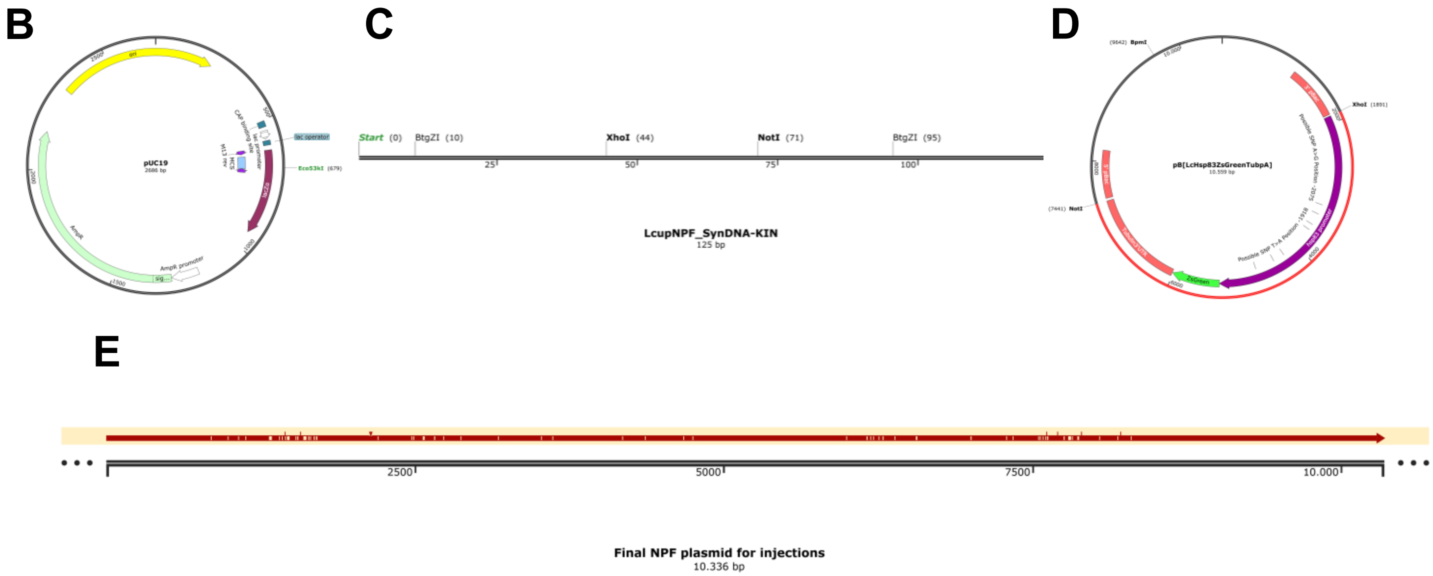


**Supplementary method 3**. *LcNPF* gene editing pipeline:

**A**. *LcNPF* gene fragment of 2042 bp corresponding to the left (L) and (R) homology arms (HAs) of the *LcNPF* homology direct repair (HDR) dsDNA donor. The fragment was obtained by PCR using the Q5® polymerase and primers (LcupNPF_K-IN_Fw-1: GTCTGTCCTCGTATACCCTCAAATA and LcupNPF_K-IN_Rv-1: ACATTACAATAACTAGGTTGCTCCA and *L. cuprina wt* gDNA obtained from adult flies of mixed sexes as template. In addition, the primer positions, restriction cutting site for BtgZI (NEB, Cat. #R0703S) which coincides with the location of LcupNPF-gRNA-11, were detailed in purple, black and red, respectively.

**B**. pUC19 vector plasmid (NEB, Cat. #N3041S) detailing the restriction cutting site Eco53KI (NEB, Cat. #NR0116S). The same plasmid was ligated with the PCR amplicon detailed in **A**, by cleaving with Eco53KI and ligated O/N at 16°C using the T4 DNA Ligase (NEB, Cat. #M0202), using an insert vector ratio of 3:1 and subsequently used for transformation of *Escherichia coli* competent cells (NEB, Cat. #C3019H) following the manufacturer's specifications. Subsequently, plasmids were obtained from 10 single clones using the ZR Miniprep Kit (Zymo Research, Cat. #D4016) following the manufacturer's specifications.

**C**. *Lucilia cuprina* NPF synthetic DNA knock-in fragment called LcupNPF_SynDNA-KIN [Integrated DNA technologies, IDT, Newark. NJ, USA, gBlocks™) detailing restriction cutting sites cutting with BtgZI, XhoI and NotI. Both the plasmid obtained in **B**, *i.e* pUC19 + *LcNPF* gene fragment, and the LcupNPF_SynDNA-KIN synthetic DNA, were cleaved with BtgZI and ligated using the T4 DNA Ligase and cloned following same protocols described before.

**D**. Donor plasmid containing the ZsGreen marker flanked by restriction enzymes XhoI and NotI (NEB, Cat. #R0146S and R3189S, respectively) (Concha et al. 2011). The plasmid obtained in **C**, *i.e*. pUC19 + LcNPF gene fragment + LcupNPF_SynDNA-KIN synthetic DNA, was cleaved using the restriction enzymes XhoI and NotI and ligated with an insert including the ZsGreen marker following same protocols described before. The insert including the ZsGreen was obtained and purified from the donor plasmid (Concha et al. 2011) previously to ligation.

**E**. The final dsDNA plasmid to be used for injections was sequenced by Oxford Nanopore (MGH CCIB DNA core, Boston, MA) and aligned to the in-silico sequence to confirm the sequence identity. For embryo microinjection, the final DNA plasmid was obtained by using a ZymoPURE II Plasmid Midiprep Kit (Zymo Research, Cat. #D4200) and further purified using the DNA Clean Kit (Zymo Research, Cat. #D4004) following manufacturer's specifications. Prior to injection the plasmid was diluted in nuclease-free (NF)-water to 1250 ng/µl concentration.

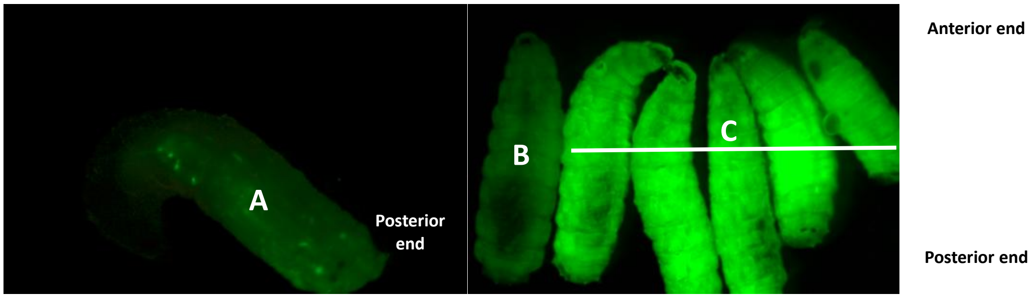

**Supplementary method 4. A**. Transient expression in mosaic *L. cuprina* larvae, and constitutive expression in heterozygous and homozygous larvae (**B** and **C** respectively) of the ZsGreen marker in seven days old L3 larvae.

**A**

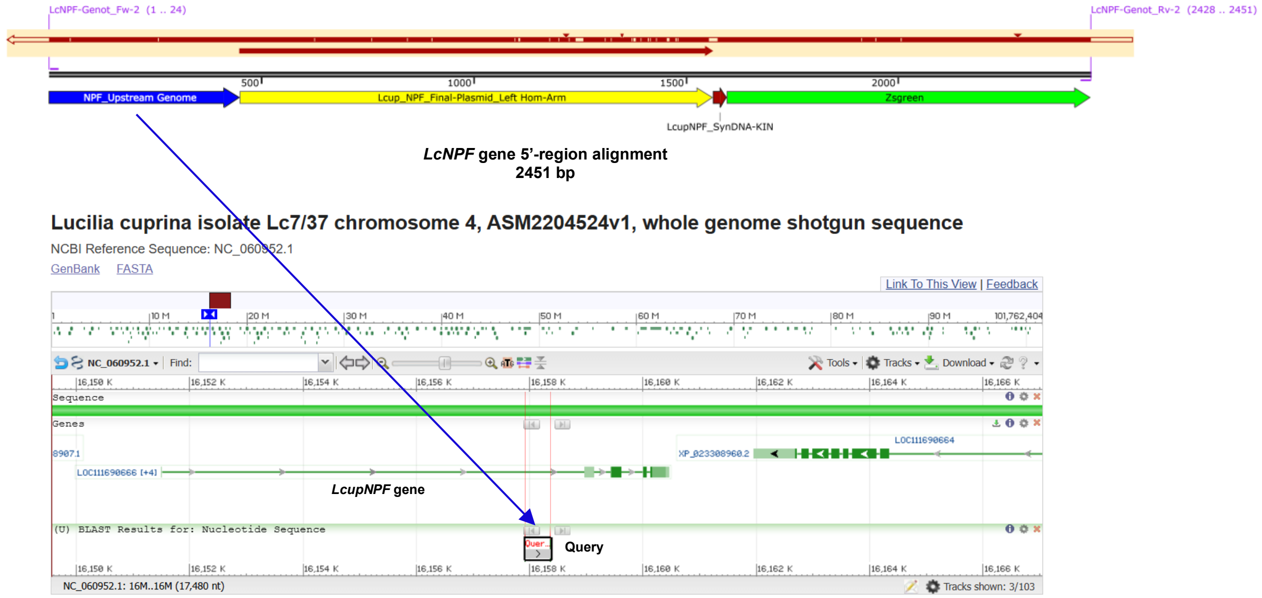


**B**

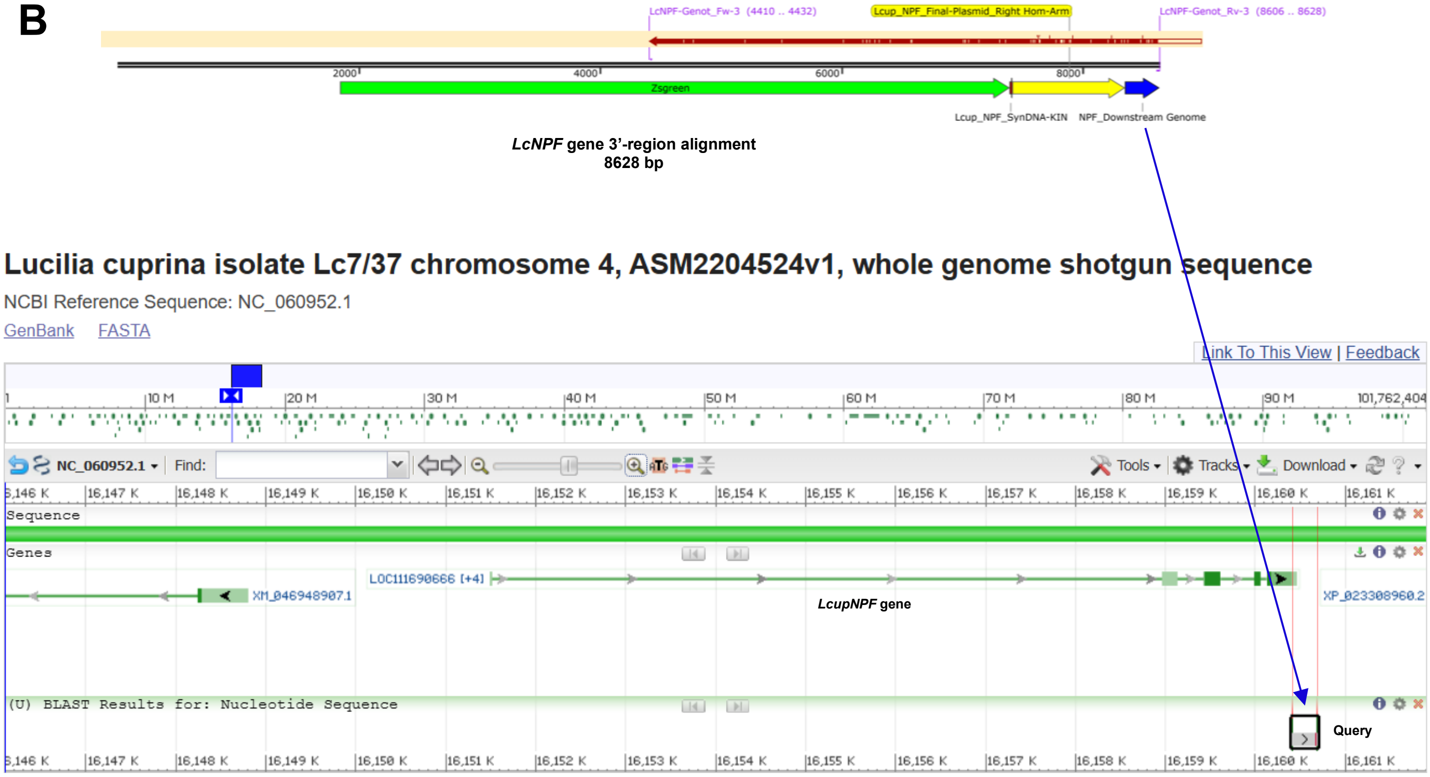


**Supplementary method 5.** Genotyping of mutated flies upstream the insert. **A**. An amplicon of 2451 bp using primers placed on the ZsGreen marker and the *LcNPF* gene (upstream the insert position), LcNPF-Genot_Fw-2: CAGCGAAACTGTTTGAAGGATCAC and LcNPF-Genot_Rv-2: CACTTCATCGAGAGGAACCATTCT respectively, and the Q5®, incubated using OneTaq® DNA Polymerase (NEB, Cat. #M0480S) to add cohesive ends and cloned using the pGEM®-T system (Promega, Fitchburg, WI, USA, Cat. #A1360), together with 10-beta competent cells (NEB), following the manufacturer's specifications. Subsequently, the plasmid was sequenced by Oxford Nanopore and aligned to the in-silico construct to confirm the sequence identity. **B**. Genotyping of mutated flies downstream the insert. An amplicon of

4218 bp using primers placed on the ZsGreen marker and the *LcNPF* gene (downstream the insert position), LcNPF-Genot_Fw-3: GGGTAGTTTGTTCCAGAACATTC and LcNPF-Genot_Rv-3: CCTACTTTGGCAAACCTGTGTTC and the Q5-Taq, incubated using OneTaq® DNA Polymerase (NEB, Cat. #M0480S) to add cohesive ends and cloned using the pGEM®-T system (Promega, Fitchburg, WI, USA, Cat. #A1360), together with 10-beta competent cells (NEB), following the manufacturer's specifications. Subsequently, the plasmid was sequenced by Oxford Nanopore and aligned to the in-silico construct to confirm the sequence identity.

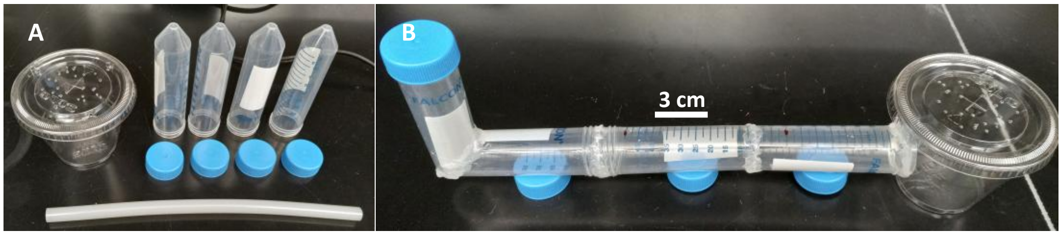


**Supplementary method 6**. Arena used for the larval locomotion assay. The approach was similar to that described by Post & Paululat (2018) except the petri dish was replaced by three 50 mL falcon tubes to improve larval crawling following a straight line. After testing late L2 and early and wandering late L3 larvae, the latter stage was selected for the assay because they showed the most consistent crawling behavior, *i.e.* crawling in a straight line for a longer time. A single larva was released at a time to assess locomotion behavior. Upon reaching the central tube, a subsequent larva was released, continuing this process until approximately 30 larvae were included per treatment. Only larvae exhibiting straight-line crawling were selected for analysis until counting 25 larvae per compared group. The time, in seconds, required for each larva to traverse 3 cm between two marks on the central tube was recorded. The starting point (timer on) was defined as the moment when the anterior tip of the larva made contact with the first mark, and the reaching time (timer off) was recorded when the anterior tip of the same larva touched the second mark. **A** and **B** show pieces used to make the area before and after assembling respectively. In addition, a short video showing the larval crawling behavior was submitted with the supplementary material.

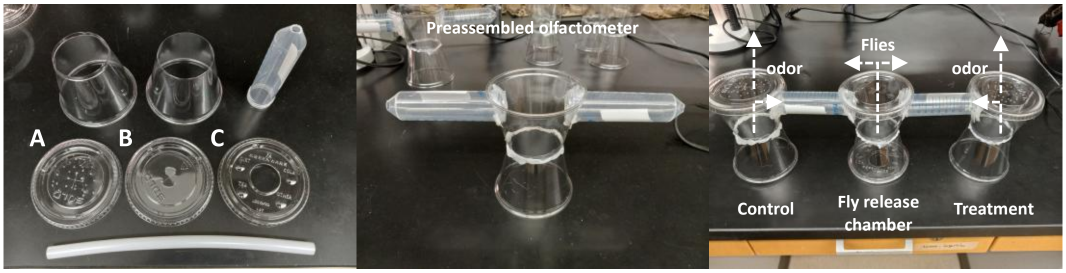


**Supplementary method 7**. Disposable static olfactometer. The static olfactometer was adapted from (Martin *et al*. 2020) and was made with disposable plastic cups, 50 mL falcon tubes (Corning®, Corning, NY, USA) and plastic glue. Flies were introduced into the central chamber, while samples were positioned in the lateral chambers. Openings of approximately 5 mm at the ends of the falcon tubes allowed the flies to access the lateral chambers and subsequently become trapped within them. Lid types: **A**. Used for top of sample chambers (small holes). **B**. Used for bottom sample chambers, and top of fly release chamber (no holes). **C**. Used for the bottom of the fly release chamber (~ 1 inch opening).

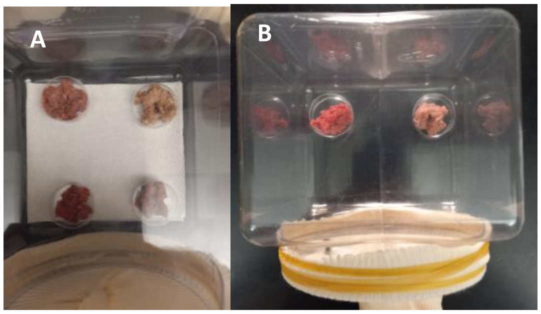

**Supplementary method 8**. Oviposition substrate preference assay. Four-inch cubic plastic bottles used for the oviposition assay showing **A** four, or **B** two oviposition substrates, respectively, as described at *Experimental procedures: Oviposition substrate preference*.
